# Structural basis for DNA targeting by the Tn7 transposon

**DOI:** 10.1101/2021.05.24.445525

**Authors:** Yao Shen, Josue Gomez-Blanco, Michael T. Petassi, Joseph E. Peters, Joaquin Ortega, Alba Guarné

## Abstract

Tn7 transposable elements are unique for their highly specific, and sometimes programmable, target-site selection mechanisms and precise insertions. All the elements in the Tn7-family utilize a AAA+ adaptor (TnsC) to coordinates target-site selection with transposase activation and prevent insertions at sites already containing a Tn7 element. Due to its multiple functions, TnsC is considered the linchpin in the Tn7 element. Here we present the high-resolution cryo-EM structure of TnsC bound to DNA using a gain-of-function variant of the protein and a DNA substrate that together recapitulate the recruitment to a specific DNA target site. We find that TnsC forms an asymmetric ring on target DNA that segregates target-site selection and transposase recruitment to opposite faces of the ring. Unlike most AAA+ ATPases, TnsC uses a DNA distortion to find the target site but does not remodel DNA to activate transposition. By recognizing pre-distorted substrates, TnsC creates a built-in regulatory mechanism where ATP-hydrolysis abolishes ring formation proximal to an existing element. This work unveils how Tn7 and Tn7-like elements determine the strict spacing between the target and integration sites.

## Introduction

DNA transposition is a widespread phenomenon that impacts many aspects of biology, from genome evolution to the spread of antibiotic resistance^1^. Transposons have evolved regulatory mechanisms to minimize the adverse effects they exert on their hosts. One of these mechanisms is target-site selection, which influences not only the impact on the host but also the future mobility of the element. The bacterial Tn7 family is known for its multiple strategies to find safe sites for insertion, ensuring the efficient spread of the element while virtually eliminating the consequences of unregulated insertion events.

The prototypical Tn7 element is the most successful and best-studied member of this family^2^. Tn7 encodes five proteins: a heterodimeric transposase (TnsA+TnsB), a molecular matchmaker (TnsC), and two distinct target-site selection proteins (TnsD/TniQ and TnsE). TnsD is a sequence-specific DNA-binding protein that targets a single *attTn7* site in the chromosome at a high frequency. TnsE is a structure-specific DNA binding protein that targets replicating structures in conjugal plasmids at low frequency, ensuring the horizontal spread of the element. In both cases, TnsC is recruited to the selected site and this, in turn, activates the transposase. Thousands of representatives on this Tn7 group have been found across diverse bacteria in different environmental niches and medically relevant pathogens^3^. Some Tn7-like elements have co-opted additional targeting proteins by fusing various DNA binding domains to the TniQ domain. In at least four independent occurrences, Tn7-like elements have co-opted CRISPR-Cas derivatives for RNA-guided transposition^4–7^. Structures of one of these CRISPR-Cas Tn7 elements revealed the interface between TniQ and the guide RNA effector complex^8^. However, the mechanism that coordinates target-site recognition with DNA integration remains unknown. TnsA, TnsB, and TnsC are conserved across the Tn7 and Tn7-like elements^6^. The TnsA+TnsB transposase has a unique cut-and-paste transposition mechanism that does not require intermediate hairpin structures. TnsB recognizes the transposon’s terminal sequences, mediating the formation of a paired donor DNA complex, and is part of the system preventing insertions at sites that already carry a Tn7 element – a phenomenon known as transposition target immunity^9^. TnsC is a AAA+ ATPase that functions as a switch for transposase activation and coordinates the interaction with the target selection proteins. TnsC recognizes TnsD/TniQ bound to the *attTn7* site or CRISPR-Cas in Tn7-like elements and directs integration with a strict spacing and orientation from the target-selection site. Similarly, TnsC also determines the orientation and spacing of insertions directed by TnsE bound to replicating structures. Gain-of-function variants of TnsC can support transposition in the absence of specific target-selection proteins, but insertions occur at lower frequencies and random sites^10^. These variants belong to one of two classes, depending on their sensitivity to target-selection proteins and whether they avoid sequences already containing a Tn7 element, highlighting the central role of TnsC in coordinating the positive and negative signals that regulate Tn7 transposition frequency.

DNA structure is also known to play an essential role in managing target-site selection. TnsD binding to the *attTn7* site introduces an asymmetric distortion on the DNA immediately upstream of the binding site^11,12^, which presumably helps recruit TnsC. Supporting this idea, the presence of a triplex DNA can drive Tn7 insertions to a hotspot located between 21-25 bp upstream to the triplex binding site when using a gain-of-function variant of TnsC. These insertions maintain the orientation bias of Tn7 insertions directed by TnsD targeting^12,13^.

Tn7 elements are functionally related to the bacteriophage Mu. Mu spreads by a replicative transposition process using the MuA transposase and the MuB regulator. In Tn7-like elements lacking TnsA, TnsB performs replicative transposition in a similar manner to MuA^14^. Moreover, TnsC is evolutionary related to the MuB regulator^15,16^. Both TnsC and MuB are AAA+ ATPases that bind DNA in an ATP-dependent, sequence-independent manner. Similar to how MuA stimulates the ATPase activity of MuB and accelerates MuB dissociation from sites nearby to another Mu element, TnsB stimulates the ATPase activity of TnsC and shuts off transposition at sites that already have a Tn7 element^9,17^. MuB forms helical filaments on DNA, and it has been proposed that the mismatch between the helical parameters of DNA and the MuB filament remodels DNA enhancing transposition^18^. The TnsABC+TnsD transposition complexes include excesses of TnsA and TnsC, supporting the idea that the oligomeric state of TnsC may be necessary for its function^19^. However, the AAA+ domain of TnsC is flanked by large extensions that mediate the interaction with the other transposon-encoded proteins (**Supplementary Fig. 1**); thus, its functional oligomeric state and DNA-binding mechanism have remained elusive.

To understand how TnsC orchestrates the interactions with other Tn7-encoded proteins and relays the presence of appropriate target sites to the TnsA+TnsB transposase, we have solved the crystal and cryo-EM structures of a gain-of-function variant of TnsC on its own and bound to DNA using a DNA substrate including an internal distortion. We find that TnsC forms heptameric open rings on DNA in the presence of a non-hydrolyzable ATP-analog. The TnsC ring has distinct N- and C-terminal faces that segregate the interactions with the target-selection proteins and the transposase to opposite faces of the ring. The N-terminal face recognizes the DNA distortion relaying the asymmetry of the loading to the recruitment of the transposase and imposing the strict spacing between the target and integration sites. This work provides key mechanistic insight to exploit the potential of programmable Tn7 elements.

## Results

### TnsC oligomerizes in a nucleotide- and DNA-dependent manner

TnsC has a central AAA+ ATPase core preceded by an N-terminal region necessary for the interaction with TnsD and followed by a C-terminal region that mediates the interactions with both TnsB and TnsA (**Supplementary Fig. 1**). TnsA interacts with the extreme C-terminus of TnsC^20^, and deletion of this region greatly improves protein stability. Therefore, we used the TnsC(Δ504-555) variant, herein referred to as TnsC short (TnsC^S^), for the structural characterization.

Most AAA+ ATPases that bind DNA do so in an ATP-dependent manner. As expected, we found that TnsC^S^ binds DNA in the presence of nucleotide in a sequence unspecific manner. A more focused shifted product was found with AMPPnP than ATP, indicating that ATP hydrolysis destabilizes the TnsC^S^-DNA complex (**Fig. 1a**). The TnsC^S^-A225V gain-of-function variant with lower ATPase activity^21^ formed a stable complex with DNA regardless of the nature of the nucleotide, confirming that ATP-binding, but not hydrolysis, is essential for the interaction with DNA (**Fig. 1a**). These results agree with previous work showing that Tn7 transposition only requires ATP binding^22^.

**Fig. 1.**
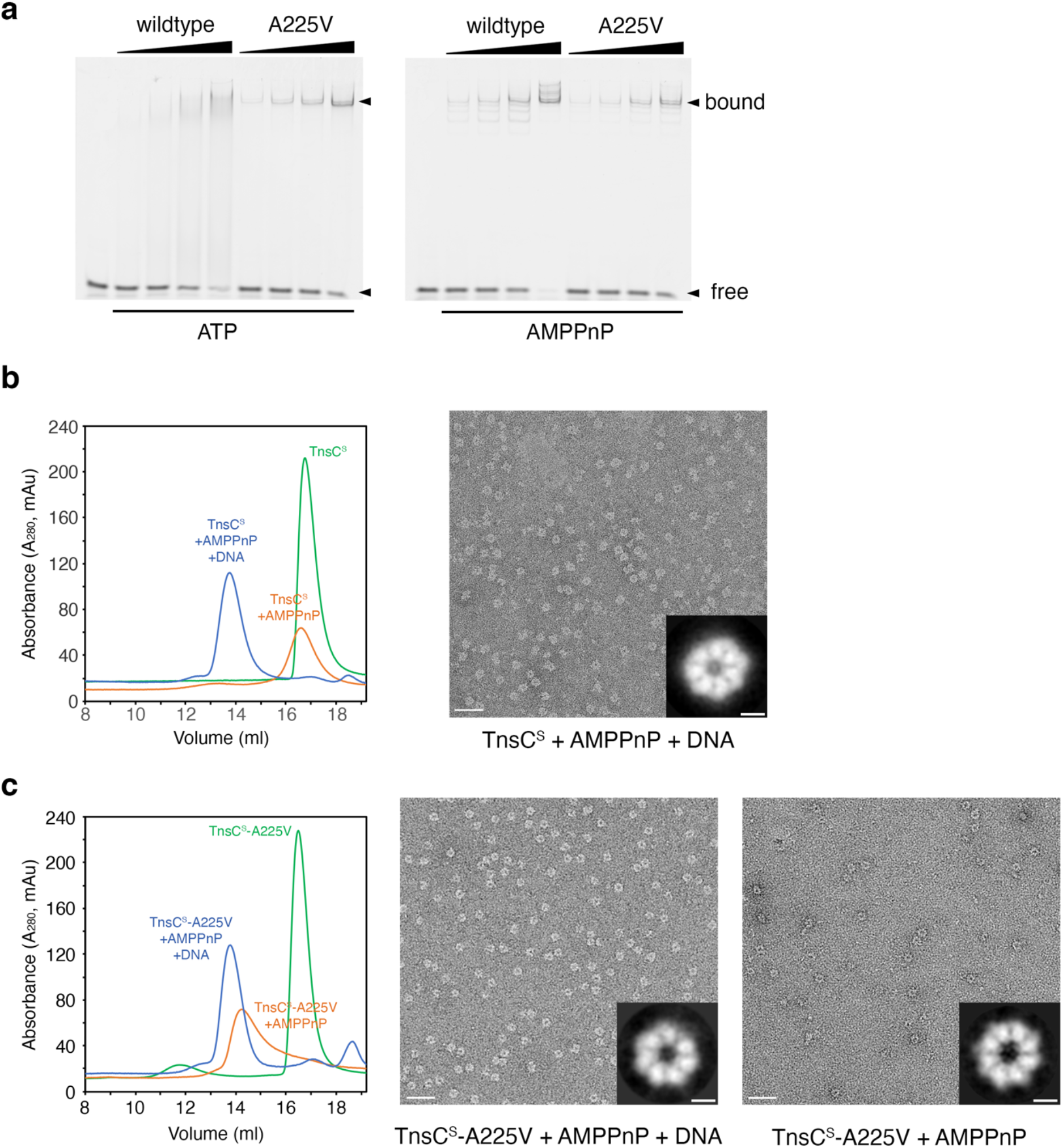
TnsC forms oligomers in a DNA and AMPPnP dependent manner. **A,** Electromobility shift assay of duplex DNA (15 bp) in the presence of either TnsC^S^ or TnsC^S^-A225V and ATP (left) or AMPPnP (right). **b,** Size exclusion chromatography profiles of TnsC^S^ on its own (green), when incubated with AMPPnP (orange), or when incubated with AMPPnP and a 15 bp DNA duplex (blue). The right-hand side panel shows a representative negative-staining micrograph and a 2D class for the TnsC^S^+AMPPnP+DNA sample. **c,** Size exclusion chromatography profiles of TnsC^S^-A225V on its own (green), when incubated with AMPPnP (orange), or when incubated with AMPPnP and a 15 bp DNA duplex (blue). The center and right-hand side panels show representative negative-staining micrographs and 2D classes for the TnsC^S^-A225V +AMPPnP+DNA and TnsC^S^-A225V+AMPPnP samples. Scale bars for the electron microscopy micrographs and the 2D classes are 50 and 5 nm, respectively.

TnsC^S^ forms large oligomers of defined molecular weight when incubated with both AMPPnP and DNA (**Fig. 1b**). TnsC^S^-A225V formed similar oligomers when incubated with both AMPPnP and DNA (**Fig. 1c**). However, TnsC^S^-A225V also formed oligomers in the absence of DNA revealing that this variant not only slows down ATP hydrolysis but it also stabilizes oligomerization. These AMPPnP-dependent oligomers seemed to be more dynamic, as illustrated by the asymmetric and tailing peak when we resolved this complex over a gel filtration column (**Fig. 1c**). Using negative staining electron microscopy, we found that both TnsC^S^ and TnsC^S^-A225V formed heptameric rings in the presence of DNA and AMPPnP, whereas TnsC^S^-A225V formed octameric rings when only AMPPnP was present (**Fig. 1b-c**).

### TnsC is a “decorated” AAA+ ATPase

We crystallized TnsC^S^-A225V and determined its structure by single-anomalous dispersion (SAD) phasing (**Table 1**). The central region of TnsC^S^-A225V (residues 74-380) adopts the characteristic AAA+ fold, but it includes three insertions (**Fig. 2 and Supplementary Fig. 1**).

**Fig. 2.**
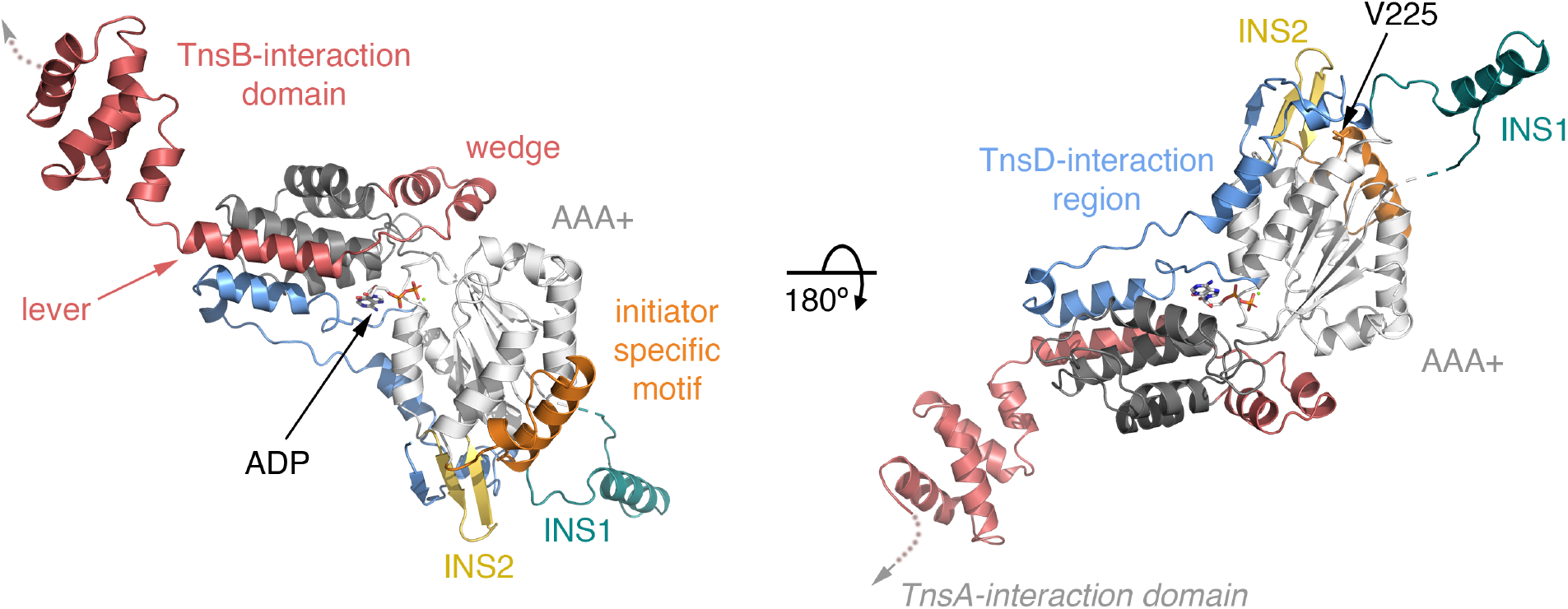
TnsC-specific insertions decorate the surface of TnsC^S^-A225V. Opposite views of the TnsC^S^-A225V crystal structure shown as color-coded ribbons: TnsD-interaction region (blue), AAA+ domain (light and dark grey for the core and lid, respectively), insertion 1 (INS1, teal), insertion 2 (INS2, yellow), initiator specific motif (ISM, orange), C-terminal extension and TnsB-interaction domain (salmon).

**Table 1.** Crystallographic data collection and refinement statistics

| **TnsC<sup>S</sup> -A225V (Sel-Met)** |  |
| --- | --- |
| **Data collection** |  |
| Beamline | CMCF 08B1-1 |
| Wavelength (Å) | 0.97856 |
| Space Group | *P*2<sub>1</sub>2<sub>1</sub>2<sub>1</sub> |
| Cell dimensions (a, b, c (Å)) | 83.78, 95.77, 313.56 |
| Resolution (Å) | 95.77-3.20 (3.32-3.20)<sup>#</sup> |
| *R*<sub>merge</sub> | 0.326 (1.418) |
| *R*<sub>meas</sub> | 0.349 (1.520) |
| *R*<sub>pim</sub> | 0.126 (0.548) |
| Completeness (%) | 100 (100) |
| Redundancy | 14.5 (14.7) |
| CC ½ (%) | 99.4 (39.3) |
| *I*/σ(*I*) | 9.7 (2.3) |
| Total reflections | 619,432 (64,911) |
| Unique reflections | 42,690 (4,414) |
| **Refinement** |  |
| Resolution (Å) | 39.20 - 3.20 (3.31 - 3.20) |
| No. reflections | 42,581 (4,166) |
| *R*<sub>work</sub> / *R*<sub>free</sub> (%) | 21.2 / 25.0 |
| No. atoms | 15,147 |
| Protein / Ligand / Water | 15,024 / 112 / 11 |
| Mean B values (Å)<sup>2</sup> | 78.8 |
| Protein / Ligand / Water | 79.0 / 57.3 / 37.5 |
| R.m.s. Deviations |  |
| Bonds length (Å) | 0.004 |
| Bonds angle (°) | 0.86 |
| Ramachandran (%) |  |
| Favored / Allowed / Outlier | 97.51 / 2.38 / 0.11 |
| Clashscore | 14.38 |
| Rotamer outliers | 0.48 |
| Rama-Z (Ramachandran plot Z-score) |  |
| Whole | -1.94 |
| Helix | -1.42 |
| Sheet | -0.19 |

<sup>#</sup>Data in the highest resolution shell is shown in parentheses

The first insertion, INS1 (residues 97-125), forms a well-defined helix connected to α0 and β1 through unstructured linkers. The helix protrudes away from the AAA+ core, and it is stabilized by crystallographic contacts. The second, INS2 (residues 154-167), introduces a β-hairpin between helix α1 and strand β2 that stabilizes the N-terminal β-strand of TnsC (β1’). The third, between helix α2 and strand β3, corresponds to the initiation specific motif (ISM, residues 195-225) characteristic of AAA+ proteins from the initiator clade^23^. This clade groups initiator/helicase-loader proteins, as well as the MuB and IstB transposition proteins. Proteins in this clade use the ISM to bind DNA, but they do so in distinct manners depending on which face of the motif engages DNA.

The N-terminal region (residues 1-73) preceding the AAA+ domain runs along the concave surface of the domain interacting with both the AAA+ core and lid, whereas the C-terminal region (residues 381-503) protrudes away from the core of the protein, giving TnsC its characteristic elongated shape (**Fig. 2b**). Within this region, we identify two motifs. The first motif, defined by helices α10 and α11, is wedged in the AAA+ domain, while the second motif (α12) leads the TnsB-binding domain (residues 426-503) away from the core of the protein. The crystal structure includes two TnsC^S^-A225V dimers in the asymmetric unit. While the AAA+ domain is identical in the four copies of TnsC^S^-A225V, INS1 and the TnsB-binding domain adopt variable orientations (**Supplementary Fig. 2a-b**). The two protomers of the TnsC^S^-A225V dimer show complementary orientations for the TnsB binding domain. Like other AAA+ proteins, the dimers further associate along one of the unit cell axes to form filaments (**Supplementary Fig. 2c**). Unexpectedly, the four TnsC^S^-A225V copies in the asymmetric unit showed well-defined electron density for an ADP molecule and a Mg^2+^ ion (**Supplementary Fig. 3**). Since ADP was not added during purification or crystallization, it may have bound to TnsC^S^-A225V during protein expression and remained bound throughout the stringent purification steps, indicating the strong affinity of TnsC for ADP.

### TnsC forms open heptameric rings on DNA

The direct interaction between TnsC and DNA is necessary to activate transposition. However, TnsC binds DNA without sequence specificity, and, therefore, gain-of-function TnsC variants direct Tn7 insertions to random sites. The group of gain-of-function variants represented by TnsC-A225V can still be recruited to specific target sites by Tn7 target-selection proteins or DNA distortions^10–12^. To mimic this specific recruitment, we assembled the TnsC^S^-A225V – DNA complex using a symmetric DNA substrate herein referred to as the 20-7-20 substrate, including seven unpaired nucleotides flanked on each side by 20 bp duplex arms (**Supplementary Fig. 4a**). Since TnsC can bind to DNA duplexes as short as 15 bp (**Fig. 1**), we reasoned that one TnsC ring would bind to each arm of the 20-7-20 DNA substrate. The complexes were assembled with excess TnsC^S^-A225V (16:1, protein:DNA), resolved over a size exclusion chromatography column, immediately deposited on grids, and flash-frozen for data collection.

The cryo-EM micrographs and resulting 2D class averages showed a mixture of top and side views and some octameric rings representing TnsC^S^-A225V molecules that were not bound to DNA (**Fig. 1 and Supplementary Fig. 4b-c**). Initial 3D classification of the particles selected from the 2D classification generated five classes (**Supplementary Fig. 5**). Class 1 represented DNA-free octameric TnsC^S^-A225V rings. Class 2 represented double-ring assemblies presumably representing one TnsC^S^-A225V ring bound to each arm of the 20-7-20 DNA substrate. The three remaining classes represented either distorted TnsC rings or incorrectly picked particles that 2D classification had not removed. We imposed a mask around the ring showing less fragmented density (bottom ring) and classified particles in class 2 in two classes (2.1.A. and 2.1.B). The cryo-EM maps obtained from these two classes refined to 4.2 and 3.5 Å resolution, respectively (**Supplementary Figs. 5-6**). The two maps were very similar and defined an open heptameric ring resembling a spring washer, with the protomers at the opening of the ring showing weaker density (**Fig. 3a**). The protomers at the opening of the ring were more disordered in class 2.1.A. Therefore, we used the map from class 2.1.B to build a model of the TnsC^S^-A225V – DNA complex (**Table 2**).

**Fig. 3.**
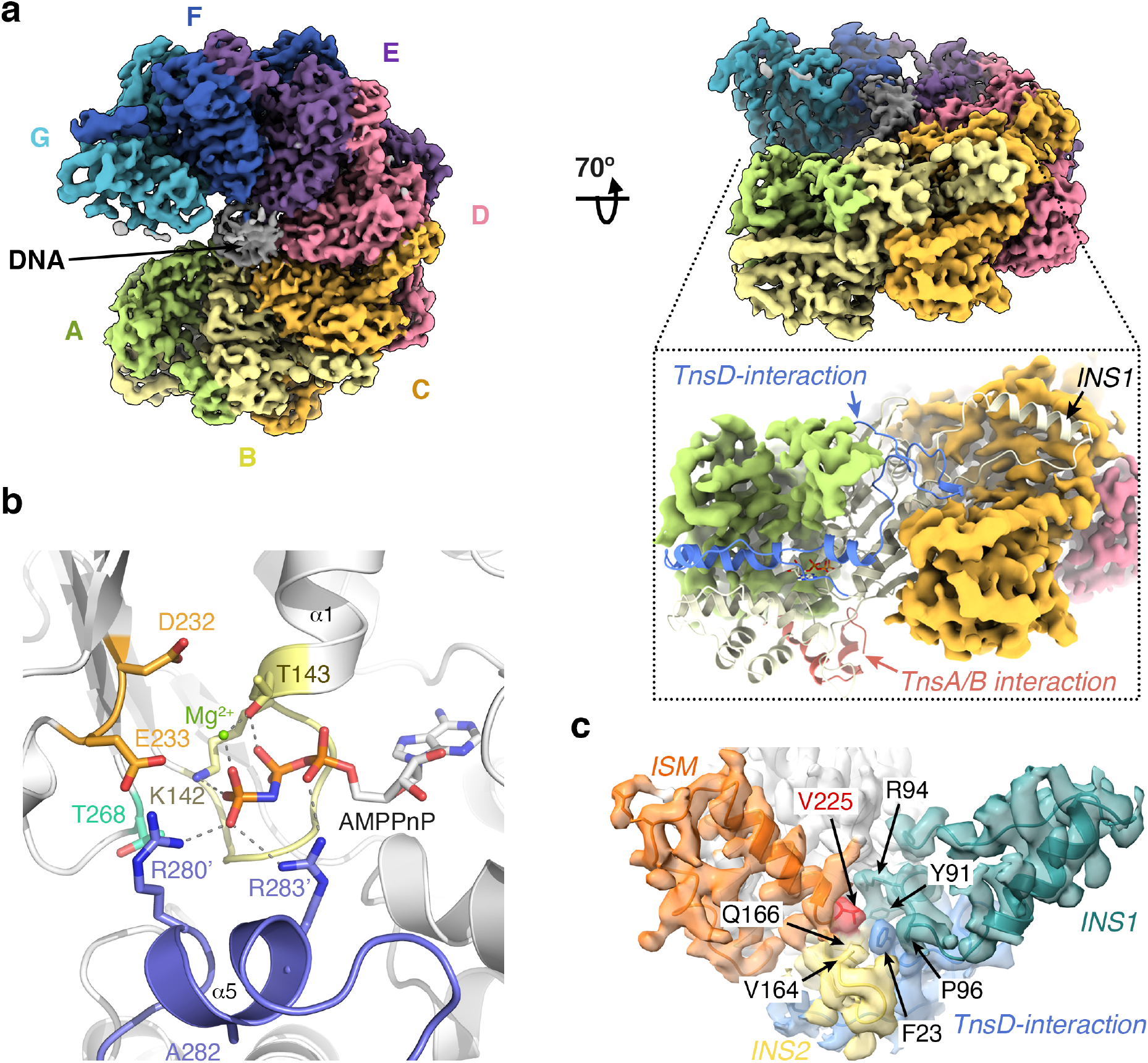
Cryo-EM structure of TnsC^S^-A225V bound to the 20-7-20 DNA. **a,** Two views of the cryo-EM map colored by subunit. The inset illustrates how the N-terminal extension preceding the AAA+ domain and INS1 extend the interface between adjacent protomers. **b**, Detail of the nucleotide-binding pocket with the conserved residues in the Walker A (yellow), Walker B (orange), sensor I (cyan), and Arg-fingers (blue) motifs shown as sticks and labeled. **c,** Detail of the cryo-EM map showing the surface pocket occupied by Val225 (red). The A225V point mutation locks the relative orientation of the three TnsC-specific insertions: INS1 (teal), INS2 (yellow), and ISM (orange). The TnsD-interaction region and the residues defining the pocket are labeled.

**Table 2.** Cryo-EM data collection, processing, and model statistics

| **TnsC<sup>s</sup> -A225V + DNA + AMPPnP** |  |
| --- | --- |
| **Data Collection** |  |
| Microscope | FEI Titan Krios |
| Voltage (kV) | 300 |
| Camera | Gatan K3 Summit |
| Defocus range (mm) | -0.5 to -2.75 |
| Pixel size (Å) | 0.855 |
| Nominal magnification | 105,000x |
| Electron dose (e-/Å<sup>2</sup>) | 78 |
| Initial particles (no.) | 905,856 |
| **Reconstruction** |  |
| Symmetry Imposed | C1 |
| Final particles | 155,988 |
| Resolution (Å) | 3.56 (post-processing)<br>3.42 (density modification) |
| FSC threshold | 0.143 |
| Map sharpening B factor (Å<sup>2</sup>) | -98 (post-processing) |
| **Refinement** |  |
| Refined model resolution (Å) | 3.4 |
| d FSC model threshold | 0.5 |
| Map CC (box/mask) | 0.64 / 0.78 |
| r.m.s.d. Bond lengths (Å) | 0.010 |
| r.m.s.d. Bond angles (°) | 1.055 |
| Molprobit score | 2.53 |
| Clash score | 14.15 |
| Rotamer Outliers (%) | 4.27 |
| Ramachandran plot (%) |  |
| Favored / Allowed / Outliers | 94.10 / 5.66 / 0.23 |
| B factors (Å<sup>2</sup>) |  |
| Protein / DNA / Ligand | 44.2 / 134.6 / 30.6 |
| Rama-Z |  |
| Whole | -3.36 |
| Helix / Sheet / Loop | -2.26 / -1.06 / -2.33 |

The refined cryo-EM map allowed the tracing of residues Ala3 to Val405 for all protomers except protomer A, where only the AAA+ core and TnsC-specific insertions were interpretable. Only six AMPPnP molecules are present in the structure because protomer A does not have an adjacent protomer to form a complete binding site. As for other multimeric AAA+ ATPases, the oligomerization interface defines a composite ATP binding sites with the Walker A (^136^GCSGSGKT^143^), Walker B (^228^LLVIDE^233^), and sensor I (T268) motifs contributed by one of the protomers and the Arg-fingers (^280^RSARR^284^) contributed by the neighboring protomer (**Fig. 3b**). TnsC lacks sensor II (D345 occupies the equivalent position), but this is not unusual in AAA+ ATPases with multiple arginine fingers^24^. The oligomerization interface of TnsC, however, extends beyond the ATP binding site. INS1 wraps around the adjacent protomer’s outer rim and interacts with the N-terminal extension (TnsD-interaction region) preceding the AAA+ domain of the adjacent protomer, covering the concave surface of TnsC^S^-A225V. Furthermore, the TnsD-interaction regions in adjacent protomers interact with each other defining the outer rim of the ring and effectively extending the buried interface from 1,631 to 2,878 Å^2^ (**Fig. 3a**). Val225 fills a small hydrophobic cavity at the base of INS1, INS2, and the initiator-specific motif (**Fig. 3c**). The presence of the bulkier side chain at this position locks the relative orientation of these TnsC-specific insertions, thereby explaining why the TnsC^S^-A225V gain-of-function variant forms rings even in the absence of DNA (**Fig. 1**).

In contrast to the crystal structure, the region encompassing helices α12-α16 was not visible for any of the cryo-EM structure protomers. The conformation of helix α12 seen in the crystal structure would preclude ring formation. This implies that the linker preceding this helix must reorient the C-terminal region of TnsC to allow for ring formation. Comparison of the crystal and cryo-EM structures reveals that Asp402 is the switch that enables ring formation and, in turn, switches transposition ‘on’ and ‘off’ (**Fig. 4**). To form the TnsC ring, Asp402 changes the direction of the main chain by about 180°, directing the C-terminal region outwards (**Movie S1**). Notably, the gain-of-function variant with the most striking increase in transposition frequency carries a mutation in this switch (TnsC-S401 YΔ402)^10^. TnsC-S401 YΔ402 represents the group of gain-of-function variants that are not sensitive to target-selection proteins, suggesting that reorienting the C-terminal region of TnsC, and consequently the TnsA and TnsB interacting domains, outside the ring may be one of the checkpoints to authorize transposition.

**Fig. 4.**
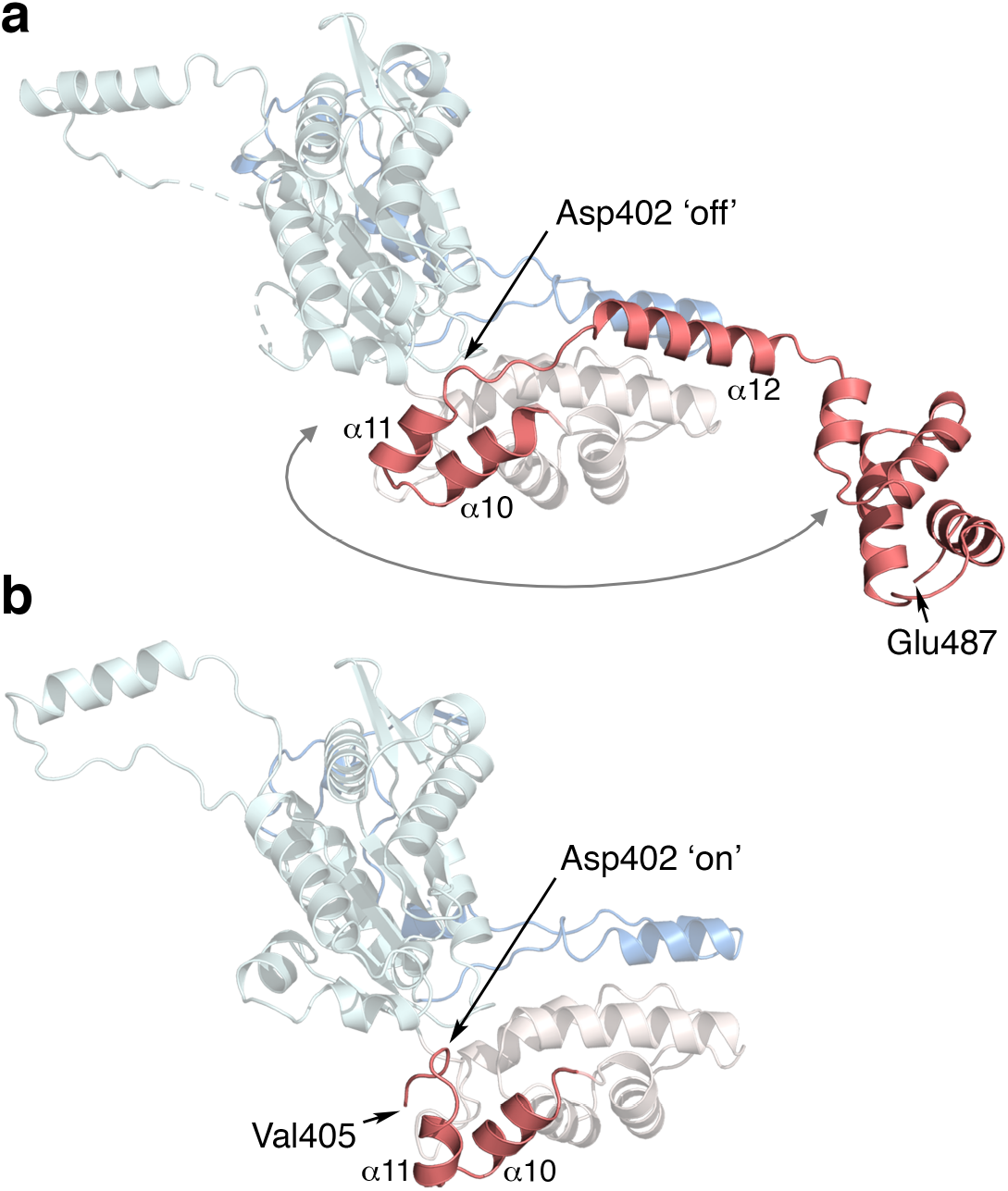
Asp402 orients the C-terminal region of TnsC. **a,** Ribbon diagram of the TnsC^S^-A225V crystal structure colored in blue (TnsD-interaction region), cyan and light pink (AAA+ domain core and lid, respectively) and coral (C-terminal extension). **b,** TnsC^S^-A225V protomer in the cryo-EM structure with the shown from the inner surface of the ring and with the same color scheme as in **(a)**. The rotation around Asp402 that the C-terminal extension must undergo to assemble the TnsC ring is indicated with a grey arrow. See also **Movie S1**.

### TnsC is a minor groove DNA binding protein

Many AAA+ ATPases from the initiator clade use the initiator-specific motif (ISM) to form helical filaments to bind, and in most cases remodel, DNA^15^. The inner surface of the TnsC ring is concave, with the central channel narrowing at the N-terminal face of the ring. DNA threads perpendicularly to the ring, with the inner rim of the N-terminal face defined by the initiator-specific motif anchoring it in place (**Fig. 5a**). As proposed from footprinting studies^11^, TnsC binds to the minor groove of the DNA duplex. Arg207, at the apex of the ISM, protrudes into the minor groove without forming any sequence-specific contacts but providing favorable electrostatics (**Fig. 5b**). Lys206 (ISM) and Lys181 (at the beginning of helix a2) interact with the phosphate backbones delimiting the minor groove (**Fig. 5b**), engaging both strands of the duplex. Point mutations in Lys206 and Arg207 weaken but do not abolish DNA binding, but the TnsC^S^-K181A/K206S/R207A variant completely loses DNA binding (**Fig. 5c**). Like the individual K206S and R207A mutations, the single K181A mutation reduced DNA binding only marginally, indicating that the interactions with the phosphate backbone of the two DNA strands are essential to anchor DNA.

**Fig. 5.**
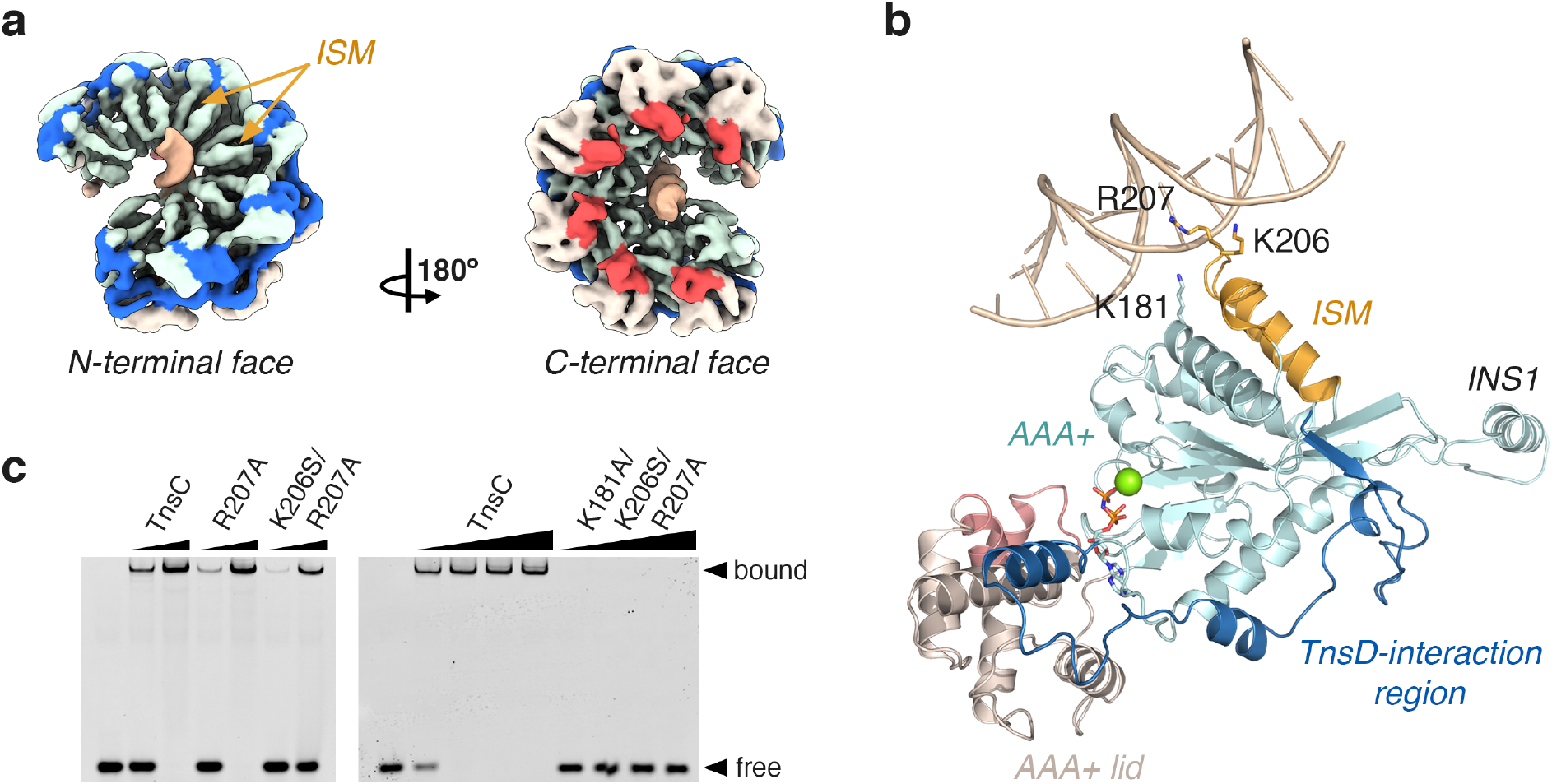
TnsC binds DNA through the initiator specific motif (ISM). **a,** N- and C-terminal faces of the TnsC^S^ ring shown in the same color scheme as in Fig. 4. **b,** Detailed view of the interactions between one TnsC^S^-A225V protomer and DNA. The ISM highlighted in gold with the residues involved in DNA-binding shown as sticks. **c,** Electromobility shift assays of a 15 bp duplex DNA (10 nM) with TnsC^S^, TnsC^S^-R207A, TnsC^S^-K206S/R207A and TnsC^S^-K181A/K206S/R207A. Protein concentrations are: 70-280 nM (left gel), and 70-560 nM (right gel).

Each pair of TnsC^S^ protomers shields three base pairs of DNA (1.5 bp/protomer), with the heptamer contacting a complete DNA turn (10.5 bp). This results in the DNA duplex retaining a B-form and revealing that TnsC does not remodel DNA. This is in contrast to most AAA+ ATPases from the initiator clade^18,23^. The central part of the TnsC ring is wider, defining a continuous solvent-accessible cavity along the major groove of the DNA. At the opposite face of the ring, the α3-α4 loop closes into the phosphate backbone of the minor groove. This α3-α4 loop includes several positively charged residues. However, they do not interact directly with the phosphate backbone, and point mutations in this loop do not affect DNA binding, indicating that this face of the ring may provide favorable electrostatics, but it is not necessary for DNA binding.

### The N-terminal face of the TnsC recognizes the DNA distortion

The fact that the TnsD-interaction region and the transposase binding domains are in opposite faces of the TnsC ring, suggests that TnsC imposes a partitioning mechanism to coordinate target-site selection and transposase recruitment. We, therefore, set to elucidate what face of the TnsC^S^ ring recognized the unpaired region of the 20-7-20 DNA substrate. To this end, we 3D classified the particles forming double rings (class 2) without applying any mask and obtained five classes (**Supplementary Figs. 5-6**). Four of these classes had two rings bound to DNA (classes 2.2.A-2.2.D in **Fig. 6a**), while the fifth (class 2.2.E) only had one ring and was not considered further. The bottom rings were well defined in all four classes, but the top ring was only clearly defined in class 2.2.A (**Fig. 6a**). In class 2.2.A, DNA showed continuous density across the entire particle with the C-terminal face of both TnsC^S^-A225V rings oriented towards the center of the particle (**Fig. 6b**). For classes 2.2.B-2.2.D, the top rings only showed fragmented densities, and the density corresponding to DNA was clear within each ring but disappeared in the center of the particles (**Fig. 6a**). This suggested that the two TnsC^S^ rings sandwiched the central bubble of the 20-7-20 substrate, with the bubble providing enough flexibility for the rings to move with respect to each other. In these three classes, the N-terminal face of the bottom ring was oriented towards the center of the particle, as illustrated by the characteristic features of the initiator specific motif (**Fig. 6b, bottom**). We then refined the top ring of the particle by focused refinement (**Figures S5-7**). In all three classes, the top TnsC^S^-A225V ring also had the N-terminal face oriented towards the center of the particle (**Fig. 6b, top**).

**Fig. 6.**
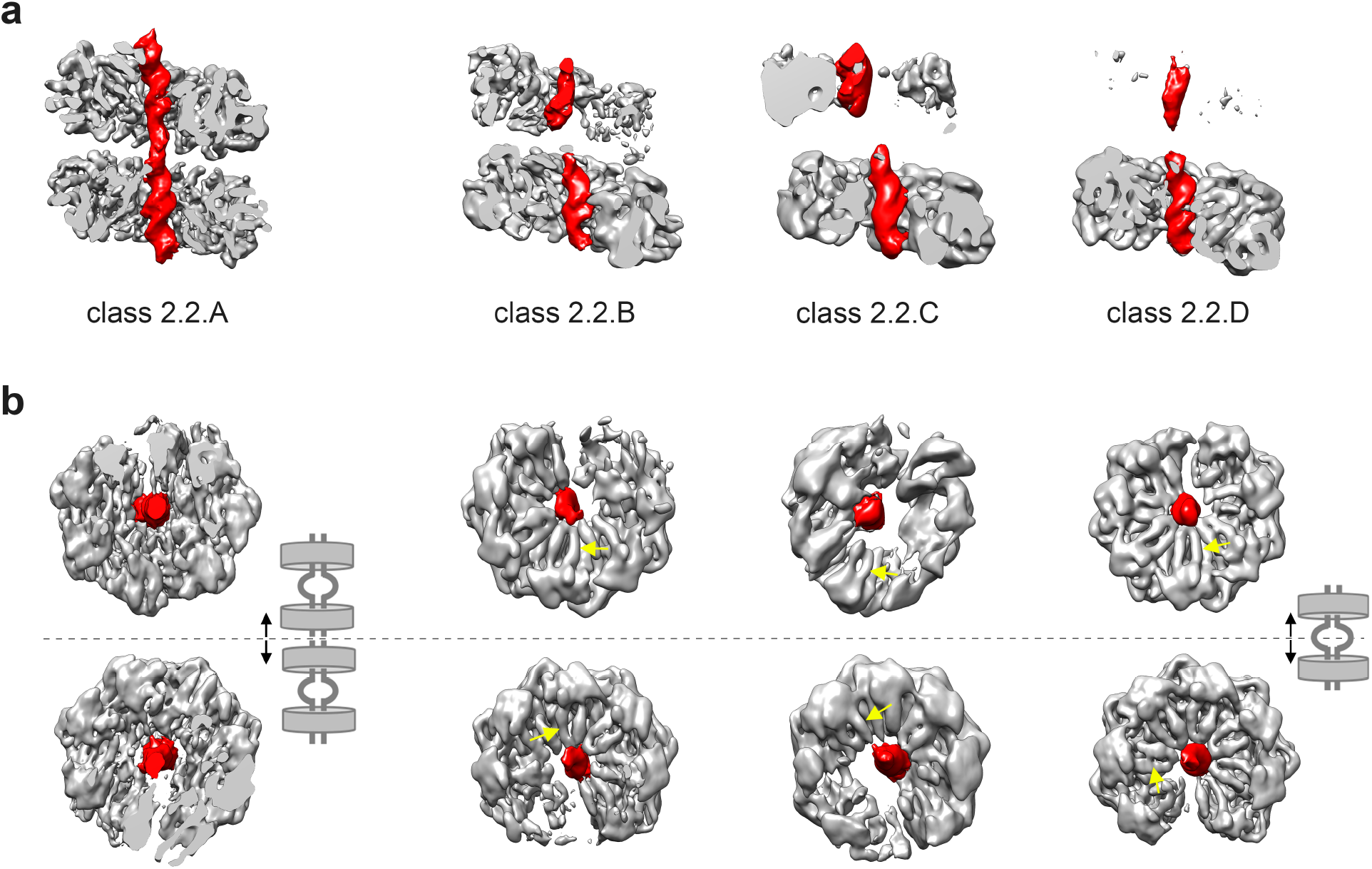
The N-terminal side of the TnsC ring faces the DNA distortion. **a,** Cross-sections of the unsharpened cryo-EM maps for classes 2.2.A – 2.2.D with densities corresponding to DNA colored in red (see Figs. S 4-6). **b,** Top (top row) and bottom (bottom row) ring for each class viewed from the center of the particle. The N-terminal face of the ring is characterized by the initiator-specific motif (yellow arrows).

This analysis revealed that the N-terminal face of the TnsC^S^-A225V ring recognizes the DNA distortion and that classes 2.2.B – 2.2.D represented particles formed by the assembly of a TnsC^S^-A225V ring on each arm of the symmetric 20-7-20 DNA substrate (**Fig. 6b**). In contrast, class 2.2.A had captured end-to-end stacking of two TnsC^S^-DNA complexes held together by a combination of protein-protein interactions and DNA stacking (**Fig. 6b**). This was not unexpected because TnsC^S^-A225V rings tend to stack end to end when assembled on either short DNA duplexes or plasmid DNA (**Supplementary Fig. 8**). TnsC rings orient randomly within these stacks, representing unspecific interactions that are likely disrupted when additional Tn7-encoded proteins are present.

### Recognition of pre-distorted target sites uncouples ATP-hydrolysis from Tn7 transposition

TnsC^S^-A225V specifically recognizes pre-distorted DNA target sites (by TnsD binding, the presence of a DNA triplex, or like in this study, a ss-ds DNA junction) to assemble a ring with a pre-determined orientation. In contrast to the adaptor factors in other transposable elements^18,23^, TnsC functions as a distortion sensor but does not remodel DNA to promote insertions. In doing so, it forgoes the ATP-hydrolysis need for Tn7 transposition^25^. Accordingly, a TnsC variant carrying a point mutation in the Walker B motif (E233K) that disrupts ATP hydrolysis without affecting ATP binding is a gain-of-function variant^10^. Interestingly, the TnsC-E233A and TnsC-E233Q variants have much higher transposition frequencies than TnsC-E233K (**Supplementary Fig. 9**). These two variants carry more conservative mutations than TnsC-E233K, indicating that while ATP-hydrolysis may not be required to activate Tn7 transposition, the structural integrity of the ATP-binding site is still important. In our cryo-EM structure, the conformation of the catalytic site is poised for hydrolysis (**Fig. 3b**), confirming that the TnsC^S^-A225V bound to the 20-7-20 DNA substrate captures the functional form of TnsC.

### The ring structure determines the strict spacing between the target and insertion sites

DNA footprinting studies have shown that TnsD protects the region +25 to +55 from the insertion site and causes an asymmetric distortion of the 5’ end binding region^11^. Similarly, Tn7 insertions directed by a triplex DNA occur 21-25 bp upstream of the triplex site^12^. Binding of TnsC extends the protected DNA region by ~18 bp, covering the region +7 to +55 from the insertion site^25^. The TnsC^S^-A225V ring covers 15bp with residual density visible as the duplex emerges from the C-terminal face of the ring. Therefore, by encircling DNA, TnsC determines the strict spacing between the target and the insertion sites.

The last hundred residues of TnsC^S^ (residues 406-503) are not visible in our cryo-EM structure, but the crystal structure includes residues 1-487. The region missing in TnsC^S^ (residues 504-555) had previously been visualized in complex with TnsA^20^, allowing us to build a model of fulllength TnsC (**Fig. 7a**). The TnsC is made up of a core defined by the heptameric ring (residues 1-406) and flexible extensions (residues 407-555) protruding from the C-terminal face of the ring. These TnsC extensions contain the binding motifs for TnsB (residues 451-494) and TnsA (residues 504-555). Their flexibility may facilitate the formation of a paired complex and transposase recruitment to insertion sites. Indeed, the region immediately preceding the TnsA-interacting region (residues 495-509) is positively charged (**Supplementary Fig. 1**) and it is important for TnsA binding to DNA^20^. These flexible tails can collapse onto the C-terminal face with the footprint of the AAA+ ring determining the minimum spacing between the DNA distortion and the integration site.

**Fig. 7.**
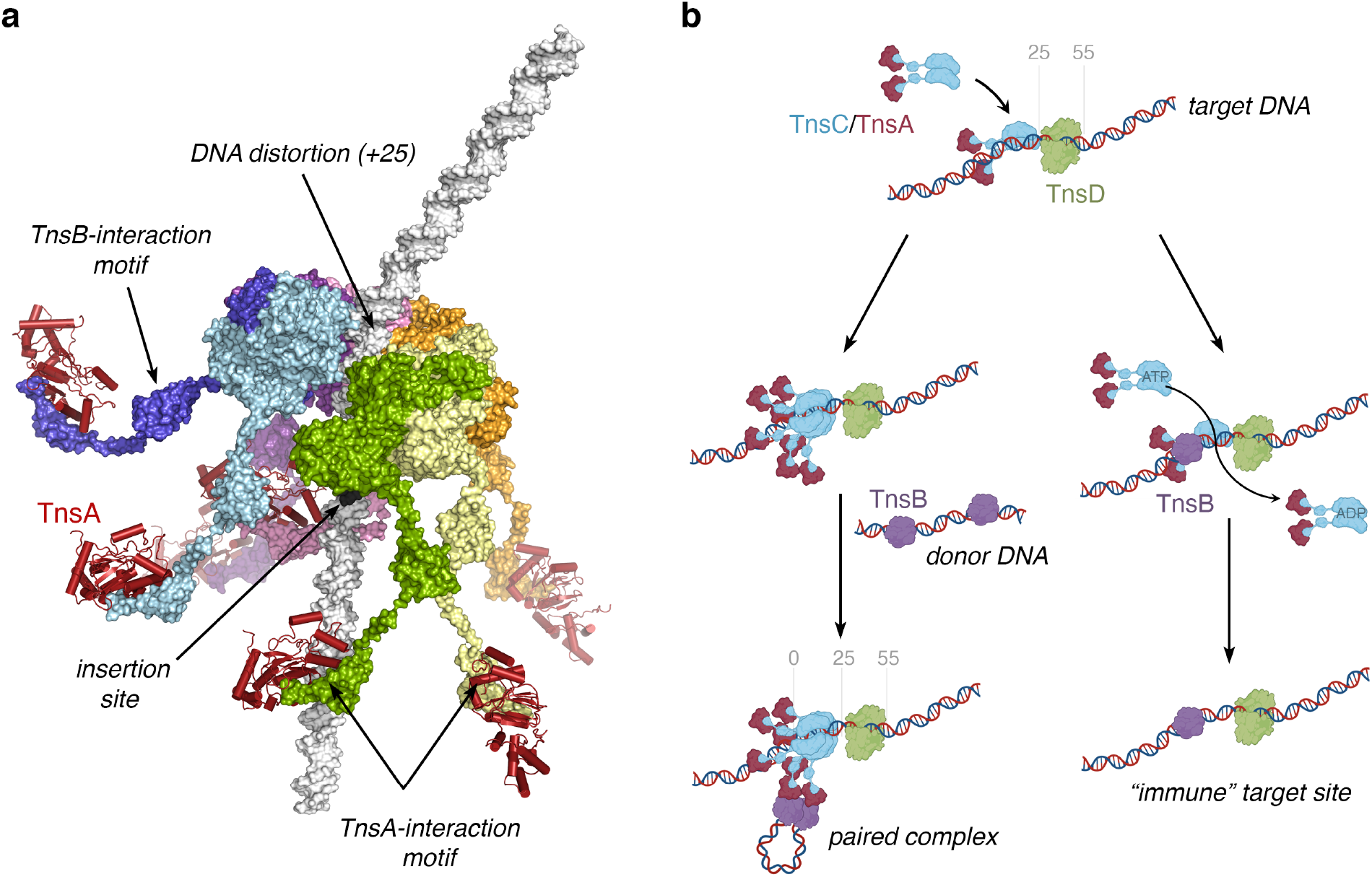
Model for DNA targeting by the Tn7 element. **a,** Model of the TnsA/TnsC complex built from the cryo-EM structure of the TnsC^S^-A225V – DNA complex (residues 1-402), the crystal structure of TnsC^S^ (residues 403-487), the crystal structure of TnsA bound the C-terminal region of TnsC (residues 506-555, PDB ID: 1T0F), and a B-DNA with a distortion at the N-terminal face of the TnsC ring. The TnsC is shown as a rainbow surface and TnsA as a dark red ribbon. **b,** Cartoon depicting Tn7 targeting and Tn7 immunity. TnsD binding to the *attTn7* site imposes an asymmetric distortion that serves as the signal for the recruitment of TnsC. TnsC can then form a ring on DNA with the N-terminal face of the ring close to the distortion and the TnsA-bound C-terminal tails protruding towards the insertion site. The TnsA/TnsC complex recruits TnsB bound to the transposon ends and stabilizes the formation of a paired DNA complex. If the site already contains a Tn7 element, TnsB can bind at the ends of the element and prevent the formation of the TnsC ring by stimulating its ATPase activity.

## Discussion

The structure of TnsC^S^-A225V reveals how a DNA distortion, presented by an ss-ds DNA junction, drives the asymmetric loading of TnsC^S^-A225V rings and imposes the strict spacing between the target and insertion sites. The structures of the TnsC dimer and the TnsC ring also explain a large body of biochemical and genetics work on the Tn7 element, and reveal how TnsC imposes the strict spacing between the target and integration sites. TnsC-A225V but not wild-type TnsC can specifically direct transposition to a DNA distortion^12^, indicating that stabilizing the TnsC ring is also an essential aspect of the target-selection process (**Fig. 1**). Binding of TnsD to the *attTn7* site induces an asymmetric distortion on the DNA^11^, but TnsD also interacts with TnsC^26,27^. It is conceivable that the interaction with TnsD stabilizes ring formation. This assigns a dual role to TnsD: marking the target site and stabilizing the TnsC ring, likely explaining why TnsD remains bound to the post-transposition complex^19^. CRISPR-Cas Tn7 elements also distort the target site with TniQ binding at one end of the distortion^8^, suggesting a similar mechanism to recruit and stabilize the TnsC ring at these sites.

The model of full-length TnsC suggests that once the ring is formed at a target site, it brings TnsA to the complex but at a specific distance from the target site (**Fig. 7a)**. TnsA molecules bound at the end of the TnsC tails can then function as beacons to recruit TnsB bound to donor DNA, enabling the formation of a paired complex (**Fig. 7b**). The uncoupling of ATP hydrolysis from recruitment of the transposase and the integration reaction presents an elegant mechanism to manage target-site selection and immunity^28^. TnsC^S^-A225V rings form in ATP or AMPPnP but not ADP, confirming that ATP binding is necessary to assemble the targeting complex. TnsB recognizes the inverted repeats at the ends of the Tn7 element and stimulates the ATPase activity of TnsC^26^. Therefore, TnsB binding to cis-acting transposition sequences near the *attTn7* site may preclude the formation of a functional targeting complex by chasing the ring away as it starts to form (**Fig. 7b**). Given its high affinity for ADP, ATP-hydrolysis would render TnsC unable to exchange ADP for ATP, effectively protecting sites already carrying a Tn7 element from additional integration events.

Our work reveals that TnsC ring stability is the underlying mechanism regulating the activation of the TnsA+TnsB transposase. Gain-of-function variants of TnsC activate untargeted transposition, but they are divided into two different groups depending on whether they activate transposition in a way that can still sense target-site selection and immunity signals (“regulated” variants represented by TnsC-A225V) or not (“de-regulated” variants like TnsC-E233K, TnsC-A282T, and TnsC-S401 YΔ402). TnsC variants with a direct effect in ATP-hydrolysis, like TnsC-E233K or TnsC-A282T (**Fig. 3b**), will be refractory to TnsB chasing away ring formation. They will also form intrinsically stable rings, rendering the stabilizing effects of TnsD/TniQ irrelevant and explaining why these TnsC gain-of-function variants cannot be targeted to specific sites. Similarly, variants like TnsC-S401YΔ402, which lacks the switch to reorient the C-terminal region and locks TnsC on the ring conformation, will also render the TnsC rings refractory to target-site selection and immunity signals. Conversely, variants that stabilize the conformation of the protomer found in the ring (like TnsC-A225V) will have a more subtle effect on stability and will still sense the stabilizing effects of TnsD, as well as destabilizing signals from TnsB.

The asymmetric loading of TnsC^S^-A225V rings on the 20-7-20 DNA substrate (**Fig. 6**), the footprinting data for the TnsC-TnsD complex^25^, and the strict spacing between the target and insertion sites matching the length of DNA covered by a single TnsC^S^-A225V ring confirm that the ring is the targeting form of TnsC. The formation of open rings also suggests a mechanism where the terminal subunit could detect the DNA distortion imposed by TnsD and reorient its C-terminal region to promote TnsC oligomerization. This mechanism would explain how dimeric target-site selection proteins interact with the TnsC heptamer (^8^ and Krishnan and Guarné, *in preparation*). As the TnsC ring assembles, the massive conformational change required to reorient the C-terminal region of each protomer added to the ring may also provide a proofreading step to check if a Tn7 insertion is already present at the site (**Movie S1**). The asymmetry of the TnsC ring will propagate to the recruitment of the transposase, providing a mechanism to impose the specific orientation for the Tn7 insertion events.

By recognizing a DNA distortion imposed by binding of a target selection factor or artificially created like in the DNA substrate used in this study, TnsC overcomes the need for a standard DNA signature while imposing a precise targeting mechanism. Instead, TnsC cedes control to the partner protein TnsD/TniQ as part of a signaling pathway with exceptional efficiency and modularity not found in any other transposition system. The division of labor between TnsC and TnsD/TniQ affords Tn7 and Tn7-like elements the plasticity to pick up a variety of DNA binding domains fused to the TniQ domain for recognizing essential genes (*glmS*, *parE*, *tRNA* genes, etc.), while continuing to direct insertions at the end of the operon with unrivaled precision. This adaptation has also allowed Tn7-like elements to co-opt CRISPR-Cas systems, where features from a TniQ-effector complex are similarly used to allow orientation control at a specific distance from the region recognized. The TnsC-TniQ interoperability may also explain how multiple elements can coexist in a single host. The foundational understanding from this work paves the way for evolution-inspired rational design or re-engineering of DNA integration systems for research and therapeutic applications.

## Methods

### Protein expression and purification

The plasmid encoding *Escherichia coli* TnsC (UniProt entry P05846) was a gift from Dr. Nancy Craig’s laboratory. All TnsC variants were generated from this plasmid by site-directed mutagenesis and verified by DNA sequencing (Génome Québec Innovation Centre). TnsC variants were produced using BL21Star(DE3) cells (Life Technologies) supplemented with a plasmid encoding rare tRNAs. Cells were grown at 37 °C in LB media to an OD_600_ ~0.7, and protein expression was induced by the addition of 0.5 mM isopropyl β-D-1-thiogalactopyranoside (IPTG) at 25 °C for 5 hours. Cell pellets were harvested by centrifugation, resuspended in lysis buffer (20 mM Tris pH 8, 0.5 M NaCl, 1.4 mM β-mercaptoethanol, 10 mM MgCl_2_, 45 mM imidazole, and 5% glycerol) supplemented with a protease inhibitor mixture (1 mM Benzamidine, 1 mM PMSF, 5 μg/ml Leupeptin, and 0.7 μg/ml Pepstatin A) and lysed by sonication. The lysate was clarified by centrifugation at 39,000 g at 4 °C for 30 minutes, and the supernatant was loaded onto a 5 ml HisTrap HP column (GE Healthcare) pre-equilibrated with lysis buffer. Contaminant proteins were eluted with 45 mM imidazole, and TnsC was eluted with 210 mM imidazole. TnsC variants were further purified by size-exclusion chromatography using a Superdex 200 Increase 10/300 GL column (GE Healthcare) equilibrated with storage buffer (20 mM Tris, pH 8.0, 0.2 M NaCl, 1.4 mM β-mercaptoethanol, 2 mM MgCl_2_, and 5% glycerol). Selenomethionine-labelled TnsC^S^-A225V was produced in B834(DE3) cells grown using the SelenoMet Medium Base plus Nutrient Mix media kit (Molecular Dimensions) supplemented with L-selenomethionine (0.04mg/ml). Protein expression and purification steps were as described above with all purification buffers containing 10 mM β-mercaptoethanol.

### DNA-binding assays

Electrophoretic mobility shift assays (EMSA) were used to assess how TnsC and TnsC-A225V changed the mobility of a 15-bp duplex DNA (formed by annealing ^5^’Fam-GCAGCCAACTAAACTA^3^’ and ^5^’TAGTTTAGTTGGCTG^3^’). Increasing concentrations of protein (0-560 nM) were incubated with 10 nM DNA in 20 mM Tris, pH 8.0, 50 mM KCl, 1 mM DTT, 1 mg/ml BSA, 5% Glycerol, 0.5 mM EDTA, and 0.33 mM ATP (or AMP-PNP) for 30 min at room temperature. Reaction mixtures (18.5 μl) were resolved on 5% Tris-Glycine native gels and imaged on a Typhoon scanner (GE Healthcare).

### Crystallization and structure determination

TnsC^S^-A225V crystals grew in 23-28% PEG 5,000 MME, 0.2M (NH_4_)_2_SO_4_, 3% glycerol, and 0.1M MES, pH 6.1-6.4. The selenomethionine-labelled crystals were grown from native protein microseeds and further optimized by macroseeding. Crystals were cryo-protected by adding 12% glycerol to the mother liquor and flash-frozen in liquid nitrogen. A complete dataset was collected at the CMCF 08B1-1 beamline of the Canadian Light Source (**Table 1**). The data were indexed and integrated using iMOSFLM^29^ and scaled with Aimless^30^. The substructure was determined using SHELXC and SHELXD^31^ at a resolution cut-off of 5.0 Å. Phase improvement and model building were done using CRANK2^32^ within the CCP4i2 package^33^. Iterative cycles of model building and refinement were done in COOT^34^ and PHENIX^35^. Coordinates and structure factors for TnsC^S^-A225V are available in the RCSB Protein Data Bank (ID: 7MBW).

### Visualization of TnsC samples by negative staining EM

Complexes of TnsC^S^ (or TnsC^S^-A225V) were prepared by mixing TnsC^S^ or TnsC^S^-A225V (21 μM), 15bp DNA (3 μM) and AMPPnP (0.5 mM) in assembly buffer (20 mM Tris pH 8, 0.2 M NaCl, 1.4 mM β-mercaptoethanol, and 0.5 mM EDTA). Mixtures were incubated at room temperature for 20 minutes and resolved over a Superose 6 Increase 10/300 GL column (GE Healthcare) pre-equilibrated with assembly buffer. The peak corresponding to the complex was immediately used for negative-stain grid preparation. TnsC^S^ complexes (5μl at 0.5 μM) were applied to 400-mesh copper grids freshly coated with a continuous layer of thin carbon and glow-discharged at 5 mA for 15 seconds. Sample excess was blotted away with filter paper (Whatman #1), and grids were stained with 1% (w/v) uranyl acetate. Excess of stain was blotted away, and the grids were dried on air. TnsC^S^ complexes were imaged on a Tecnai F20 electron microscope operated at 200 kV (University of Toronto) using a room temperature side entry holder. Images were recorded using a Gatan K2 Summit direct detector camera as 30-frame movies at a nominal magnification of 25,000×. Movies had a calibrated pixel size of 1.45 Å. TnsC^S^-A225V complexes were imaged on a Tecnai F20 electron microscope operated at 200 kV at the McGill Facility for Electron Microscopy Research (FEMR, McGill) using a room temperature side entry holder. Images were collected in a Gatan Ultrascan 4000 4 k × 4 k CCD Camera System at a nominal magnification 620,000x (calibrated pixel size of 1.83 Å). For all samples, the total electron dose per image was ~ 30 electrons/Å^2^. Images were collected using a defocus of approximately −3 μm. For each sample, 8,000 ~ 10,000 particles were selected and used to generate reference-free class 2D averages using Relion 3.1^36,37^.

### Cryo-EM sample preparation and data collection

The TnsC^S^-A225V – DNA complex was prepared using a DNA substrate including 20 bp duplex arms connected by 7 unpaired nucleotides (nt). The substrate was prepared by annealing two 47-nt long oligonucleotides (YS028: 5’GGGCAGACAACGTGGCGCGTTTTTTTTTGGTCCTAGCACAGCGTATG3’ and YS029: 5’CATACGCTGTGCTAGGACCACCCCCCCACGCGCCACGT TGTCTGCCC3’). The complex was assembled in 120 μl reactions including TnsC^S^-A225V (28 μM), 20-7-20 DNA (1.75 μM) and AMPPnP (0.5 mM) in assembly buffer (20 mM Tris pH 8, 0.2 M NaCl, 1.4 mM β-mercaptoethanol, and 0.5 mM EDTA). Reactions were incubated overnight at 4°C and resolved over a Superose 6 Increase 10/300 GL column (GE Healthcare) preequilibrated with assembly buffer. The peak corresponding to the complex (3.5 μl at 15 μM) was immediately applied to holey carbon grids (c-flat CF-2/2-2C-T), and sample vitrification was done using an FEI Vitrobot Mark IV. Initial screening and optimization of conditions were performed on an FEI Tecnai G2 F20 operated at 200 kV at FEMR (McGill) using a Gatan 626 single tilt cryo-holder. Screening images were recorded in a Gatan Ultrascan 4000 4k x 4k CCD camera. Once optimal conditions were found, a data set was collected at FEMR using SerialEM software in a Titan Krios microscope operated at 300 kV. Images were recorded with a Gatan K3 direct electron detector as 30-frame movies using 3-second exposures in counting mode and a total dose per movie of 78 e^-^/A^2^. We collected the images using a defocus range of −0.5 to −2.75 μm and a nominal magnification of 105,000x, generating images with a calibrated pixel size of 0.855 Å.

### Image processing

Movies were corrected for beam-induced motion using MotionCor2^38^ or Relion’s implementation of the MotionCor2 algorithm^37^. Their CTF parameters were estimated using the Gctf program^39^. Subsequent data processing steps were done in Relion v3.1^36,37^. Particle images were selected and extracted from the micrographs using auto-picking. Extracted particles were subjected to one cycle of reference-free 2D classification to remove incorrectly picked or damaged particles. This resulted in a clean dataset comprised of 905,856 particles. These particles were subjected to a first round of 3D classification, followed by a dual refinement strategy (**Supplementary Figs. 5-7).** The initial 3D reference used for this first round of 3D classification was generated de novo using the Stochastic Gradient Descent routine as implemented in Relion. In the subsequent layers of 3D classification done for class 2, we used the resulting cryo-EM map obtained for this class at the end of the first layer of 3D classification. The 2D and 3D classifications were done using particle images binned by 2. However, we used full-size images in the refinement steps. To obtain classes 2.1.A and 2.1.B, we imposed a mask around the ring showing the most complete density (bottom ring) and classified the particles into two classes. These two classes were subjected to Bayesian polishing, CTF refinement, and a final round of 3D refinement. The refined cryo-EM map and associated model for class 2.1.B have been deposited in the EMDB (ID: 7MCS).

To obtain classes 2.2.A-2.2.E, we performed an additional 3D classification step without applying any mask (**Supplementary Fig. 5**). In these classes, the refinement strategy did not include CTF refinement or Bayesian polishing. All classes showed well-defined density for the bottom ring, but the top ring for three of the classes (2.2.B-2.2.D) was not clearly defined. To refine the top rings, we applied a mask to focus the refinement on these rings, and alignment calculations were done using stacks of particles in which the density of the bottom ring had been subtracted (**Supplementary Fig. 7**). Refinements of the bottom rings were done by applying a mask to these rings, and particle orientation was done using the standard particle stacks. Sharpening of the final cryo-EM maps and local resolution analysis was done with Relion. The average resolution for each cryo-EM map was estimated by Fourier shell correlation using a threshold value of 0.143 (**Supplementary Fig. 6**).

### Model building and refinement

Refined cryo-EM maps were further improved by density modification in PHENIX^35,40^. The monomer structure of TnsC^S^-A225V obtained from X-ray crystallography at 3.2 Å resolution was manually fit into each of the subunits of the cryo-EM map using Chimera^41^, and an ideal B-form DNA was placed in the density as a starting point. The initial model was then subjected to rounds of model building in Coot^34,42^ and global real-space refinement in PHENIX^43^. The final model was validated with statistics from MolProbity^44^. Cryo-EM data collection, reconstruction, and model statistics are summarized in **Table 2**. Figures were prepared using Pymol (The PyMOL Molecular Graphics System, Version 2.4.0, Schrödinger, LLC), UCSF Chimera^41^, and UCSF Chimera X^45^.

### Lambda hop assay

Lambda donor transposition assays were carried out as previously described^46^. Assays were performed in strain BW27783^47^. Overnight cultures grown in LB + 30 μg/mL chloramphenicol of strains with pCW15^48^ or mutant derivatives produced by site-directed mutagenesis were subcultured into the same media and grown to mid-log at 37°C. Cells were harvested and resuspended in 20 mM MgSO_4_ to a density of 1.6 x 10^9^ CFU/mL. 1.6 x 10^8^ cells were mixed with 1.6 x 10^7^ prepared λKK1 (MOI – 0.1) and incubated for 15 minutes at 37 °C. LB + 10 mM sodium citrate was added to 1 mL, and cultures were incubated with shaking for 60 minutes at 37°C before plating onto LB + 100 μg/mL kanamycin + 20 mM Na Citrate. Plates were incubated for 16-20 hours at 37°C before colony counts were made.

## Acknowledgments

We thank Dr. Nancy Craig for critical reading of the manuscript, and Dr. John Rubinstein (SickKids, Toronto) for access to his Tecnai F20 during the early stages of this project and FEMR personnel at the McGill for data collection assistance. This work was funded by the Canadian Institutes of Health Research (PJT-155941 to A.G.) and the National Institutes of Health (R01GM129118 to J.E.P.).

## Author contributions

Y.S. and A.G. conceived the study and interpreted the data. Y.S. characterized the TnsC complexes, collected data, determined and analyzed the crystal and cryo-EM structures. J.G.B. and J.O. assisted with cryo-EM data collection and processing. M.T.P. and J.E.P. performed and interpreted the lambda hop assays. A.G. and J.E.P. obtained funding for this study. A.G. prepared the manuscript with input from all authors.

## Competing interests

Cornell University has filed patent applications with J.E.P. as inventor involving CRISPR-Cas systems associated with transposons that is not directly related to this work. The rest of the authors declare no competing interests.

