## Supplemental Figures 1-9 for "Structural basis for DNA targeting by the Tn7 transposon"

[1]Department of Biochemistry, McGill University, Montreal (QC), Canada. [2]Centre de Recherche and Biologie Structurale, McGill University, Montreal (QC), Canada. [3]Department of Microbiology, Cornell University, Ithaca (NY), USA. [4]Department of Anatomy and Cell Biology, McGill University Montreal (QC), Canada. [5]Lead Contact.

**Supplementary Material:**

**Supplementary Fig. 1:** Sequence alignment of TnsC proteins

**Supplementary Fig. 2:** Crystal Packing of TnsC$^{S}$-A225V

**Supplementary Fig. 3:** TnsC$^{S}$-A225V nucleotide binding pocket

**Supplementary Fig. 4:** Cryo-EM images of TnsC$^{S}$-A225V bound to DNA

**Supplementary Fig. 5:** 3D classification and refinement workflow for the TnsC$^{S}$-A225V–DNA complex

**Supplementary Fig. 6:** Resolution analysis of the cryo-EM maps obtained for the TnsC$^{S}$-A225V:DNA complex

**Supplementary Fig. 7:** Refinement strategy for the top ring in classes 2.2.B, 2.2.C, and 2.2.D

**Supplementary Fig. 8:** TnsC$^{S}$-A225V rings stack end-to-end when bound to duplex DNA

**Supplementary Fig. 9:** TnsABC transposition frequency with TnsC gain-of-function mutants

**Supplementary Movie 1:** Conformational rearrangements of the C-terminal region of TnsC$^{S}$-A225V

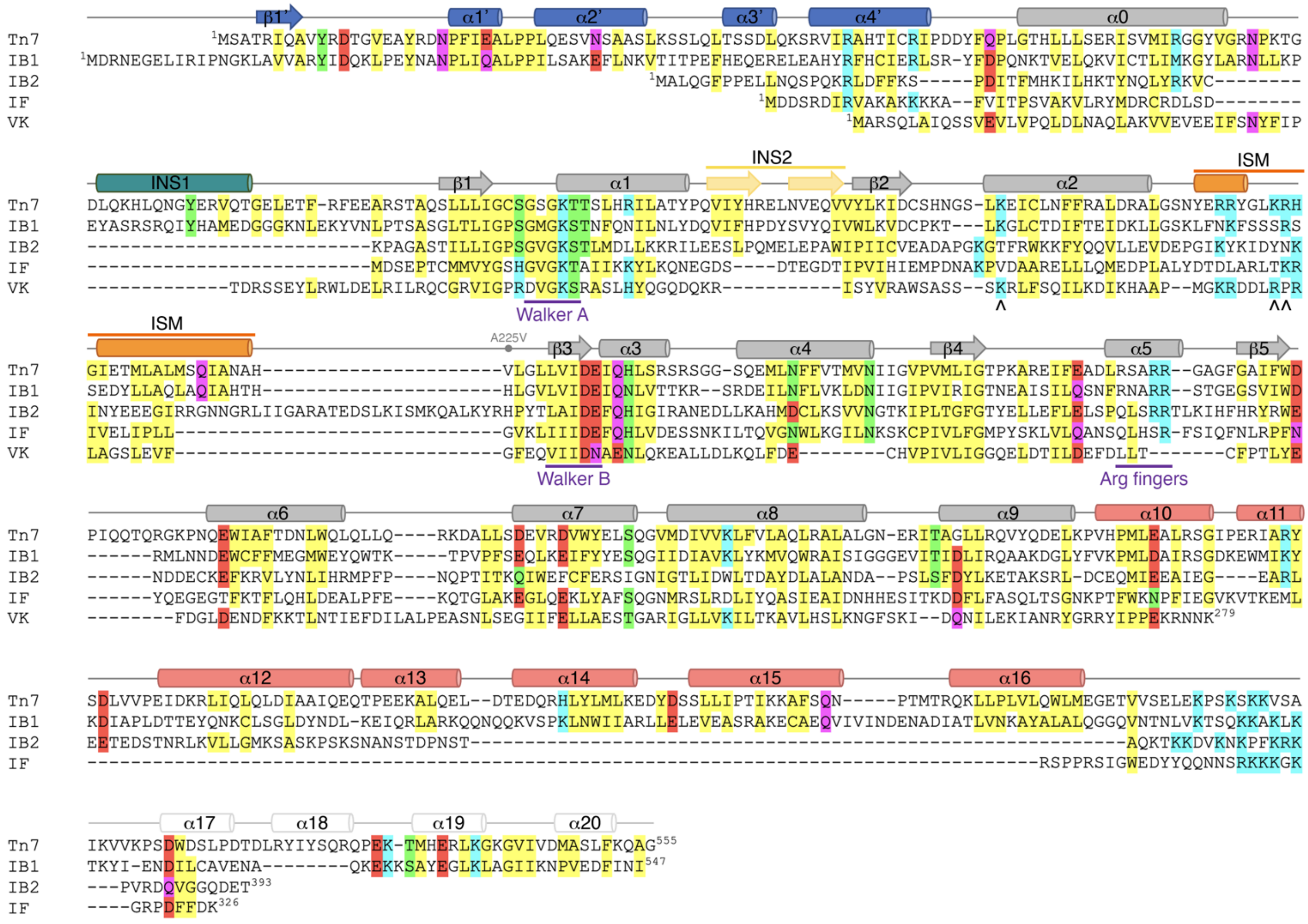


**Supplementary Fig. 1. Sequence alignment of TnsC proteins.** Alignment of representative TnsC proteins from the Tn7 element (**Tn7**: *Escherichia coli* TnsC) and Tn7-like elements including class I-B CRISPR systems (subtype 1 represented by *Nostoc calcicola* TnsC (labeled **IB1**); and subtype 2 represented by *Gloeothece verrucosa* TnsC (labeled **IB2**)), class I-F CRISPR systems (represented by *Vibrio parahaemolyticus* TnsC (labeled **IF**)), and class V-K CRISPR systems (represented by *Calothrix parietina* TnsC (labeled **VK**)). TnsC from class I-B subtype 1 is the closest to TnsC from the prototypical Tn7 element, whereas TnsC from class V-K is much shorter and lacks the characteristic N- and C-terminal extensions. Conserved hydrophobic (yellow), polar (green, pink), and charged ((-) red, (+) blue) are highlighted. Secondary structure elements from *E. coli* TnsC are shown and color-coded as in Fig. 2, and the A225V point-mutation labeled. Positively charged residues within the initiation specific motif (ISM) and involved in DNA binding are indicated with purple arrowheads.

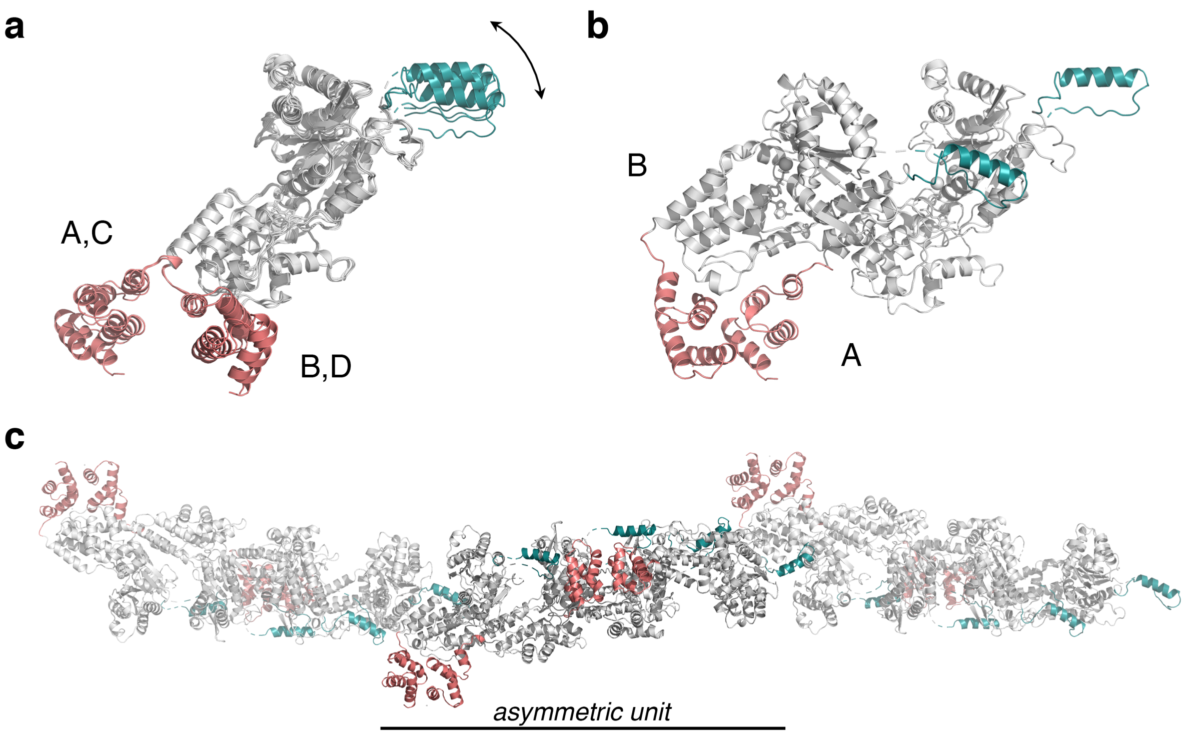


**Supplementary Fig. 2. Crystal Packing of TnsC$^{S}$-A225V. a,** Superimposition of the core region (residues 3-423) for the four molecules in the asymmetric unit returns r.m.s.d. of 0.9-1.1 Å and reveals that both the INS1 (teal) and the TnsB-interaction domain (salmon) can move with respect to the AAA+ domain. **b,** The four molecules in the asymmetric unit form two TnsC dimers (AB and CD), where each protomer of the dimer has a different orientation of the TnsB-interaction domain. The TnsC dimer interface occludes 2,200 Å$^2$, while the interaction between adjacent dimers protects 1,400 Å$^2$. **c,** In the crystal, TnsC$^{S}$-A225V dimers associate head-to-tail and define a pseudo-filament.

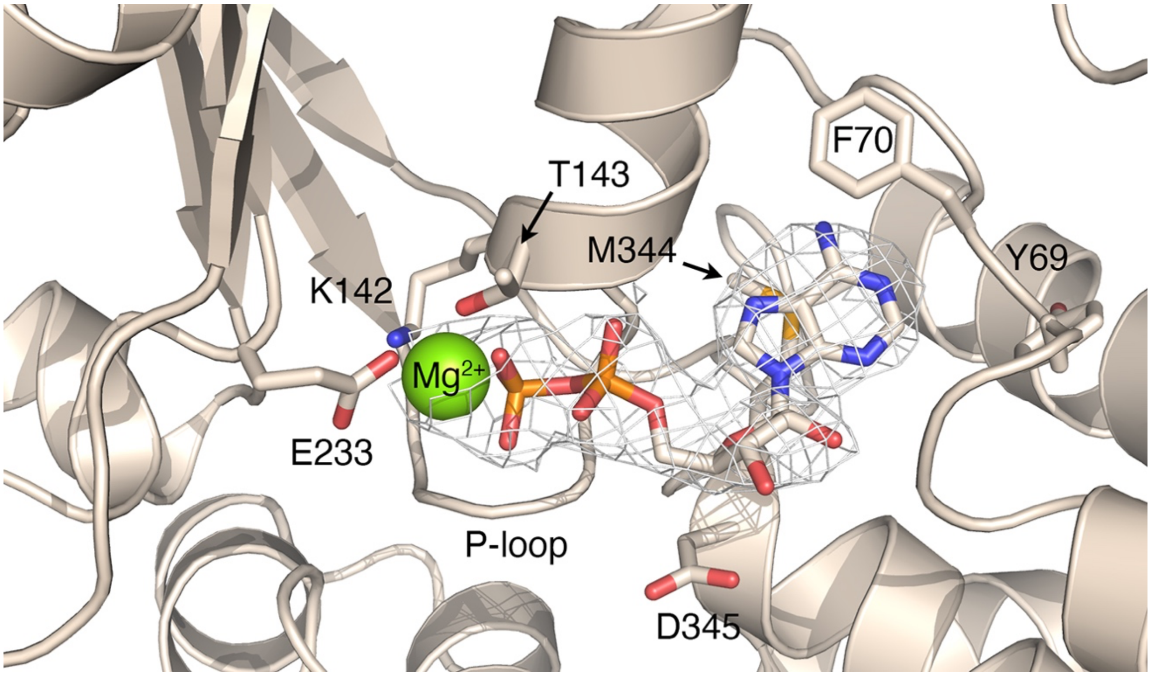


**Supplementary Fig. 3. Nucleotide-binding pocket.** Representative 2Fo-Fc electron density map (1.2 s) of the ADP molecule bound at the nucleotide-binding site of $TnsC^{S}$-A225V. All molecules in the asymmetric unit show excellent electron density, indicating the strong affinity of $TnsC^{S}$-A225V for this nucleotide. Residues in the Walker A (K142-T143) and Walker B (E233) are labeled.

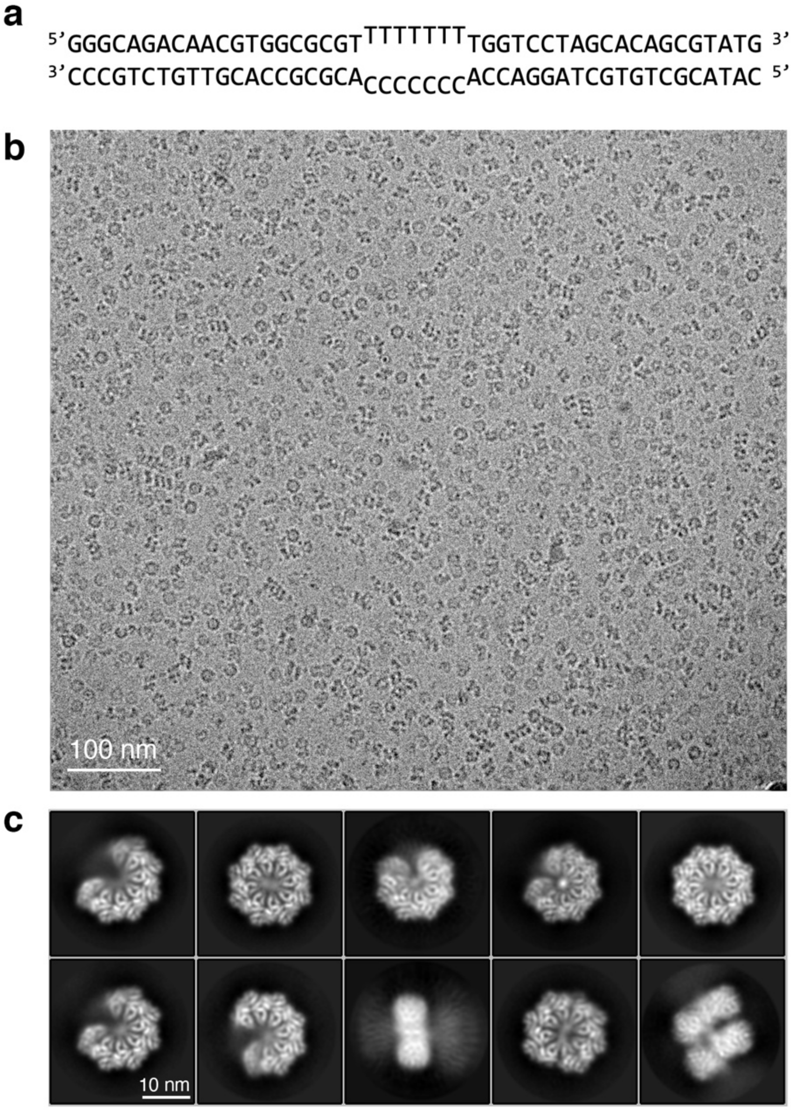


**Supplementary Fig. 4. Cryo-EM images of TnsC$^{S}$-A225V bound to DNA. a,** The sequence of the 20-7-20 DNA. **b,** Representative cryo-EM micrograph of TnsC$^{S}$-A225V bound to the 20-7-20 DNA substrate. **c,** 2D class averages show a mixture of top and side views of the complex and particles with (heptameric open rings) and without DNA (octameric closed rings).

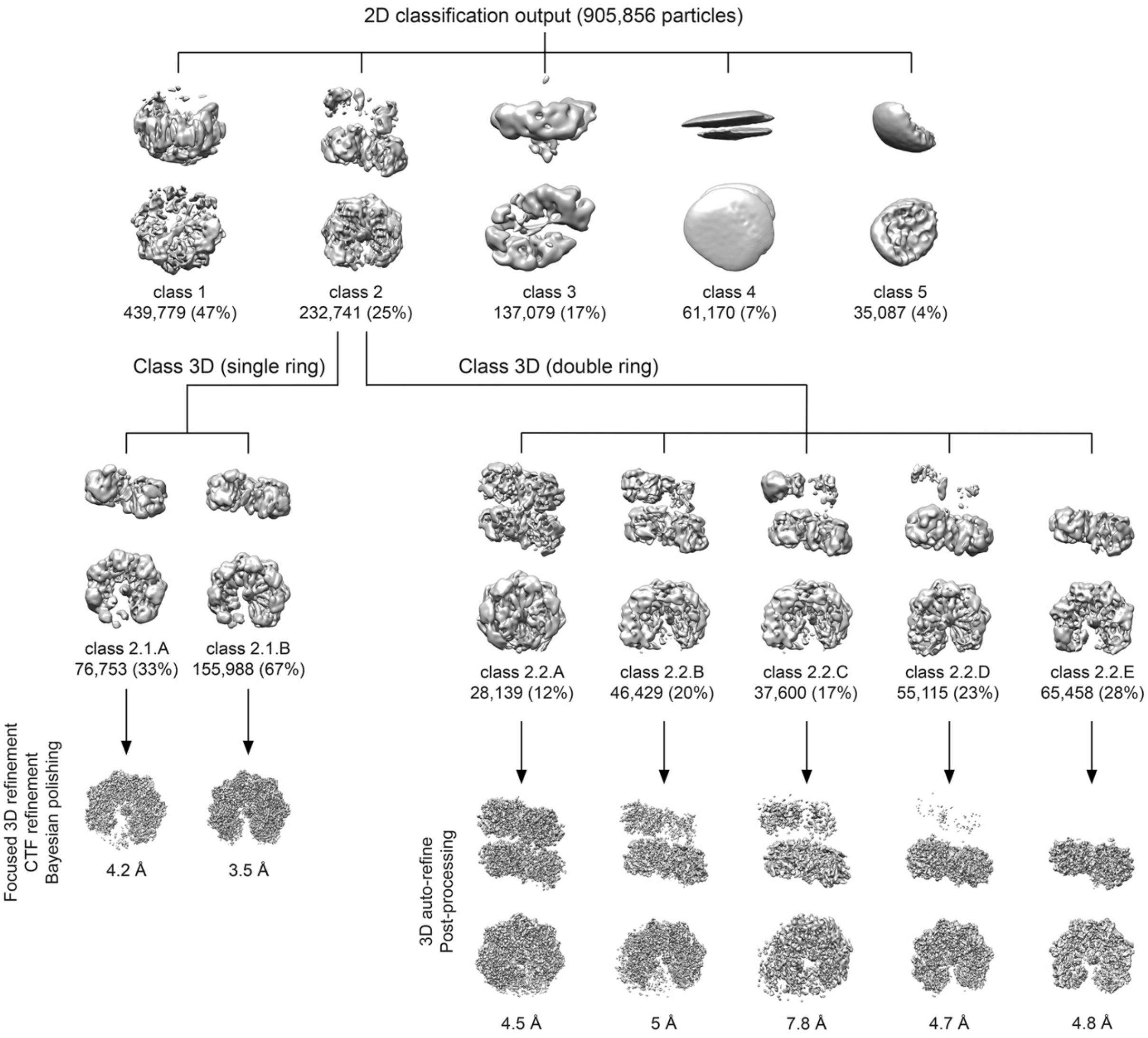


**Supplementary Fig. 5. 3D classification and refinement workflow for the TnsC$^{S}$-A225V–DNA complex.** Particle images were subjected to an initial round of 3D classification. The class, including TnsC$^{S}$-A225V particles bound to DNA (class 2), was further processed in two different ways. In a first approach (left), we imposed a mask around the ring showing the most complete density and classified the particles into two classes. Refined maps and average resolution for each of these two classes are shown on the bottom row. On a second approach (right), we 3D classified particles in class 2 without applying any mask, and we obtained five classes. Refined maps and average resolution for each of the resulting cryo-EM maps are shown on the bottom row. Percentages for each class are calculated within each subgroup.

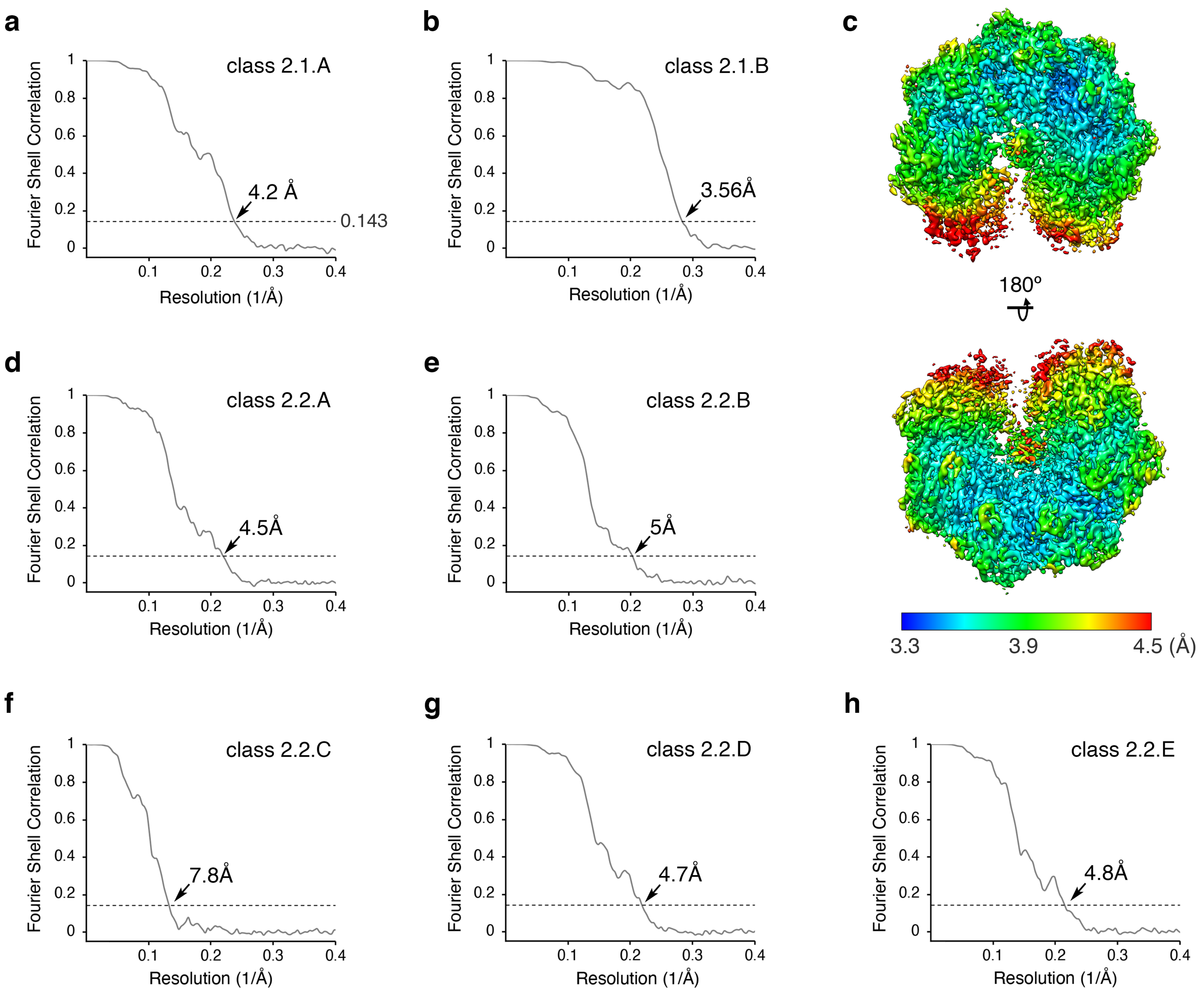


**Supplementary Fig. 6. Resolution analysis of the cryo-EM maps obtained for the TnsC$^{S}$-A225V:DNA complex.** Fourier shell correlation (FSC) plots for the final map for classes 2.1.A **(a)** and 2.1.B **(b)**. Overall resolution of the cryo-EM maps was determined using an FSC threshold value of 0.143. **c,** Local resolution analysis of the final cryo-EM map for class 2.1.B. **d-h,** Fourier shell correlation (FSC) plots for the final maps for classes 2.2.A-2.2.E.

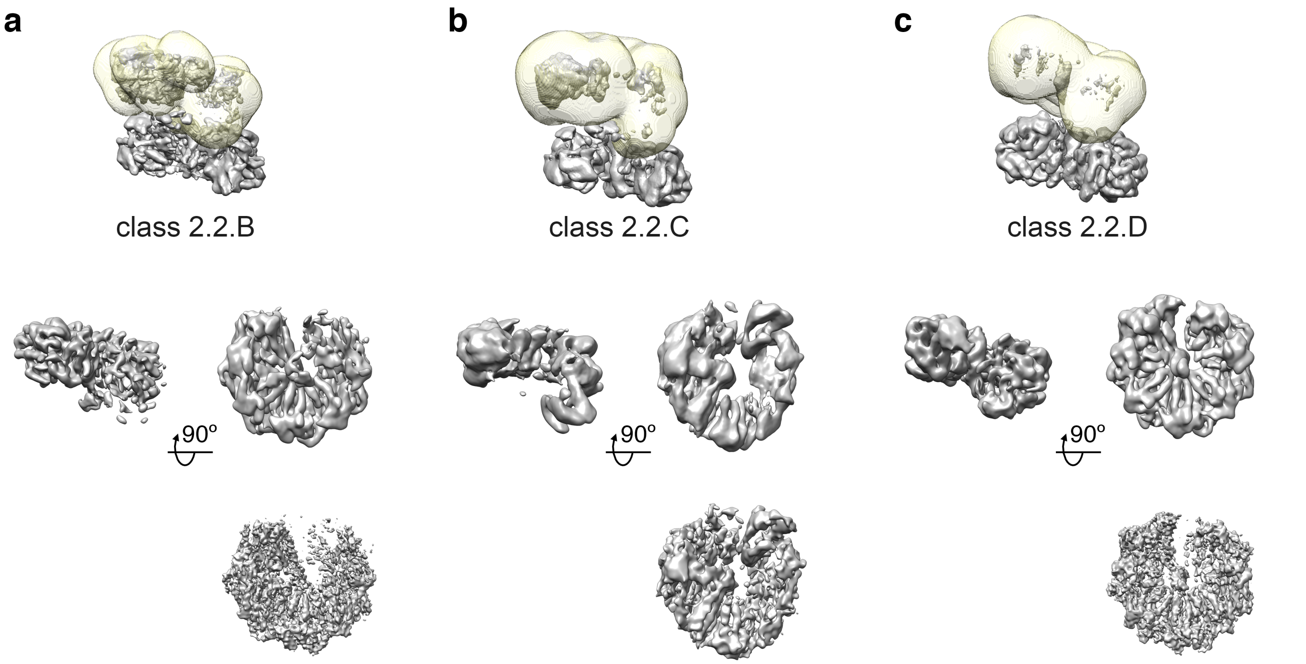


**Supplementary Fig. 7. Refinement strategy for the top ring in classes 2.2.B, 2.2.C, and 2.2.D.** For each class (**a –** class 2.2.B, **b –** class 2.2.C, **c –** class 2.2.D), refinements were performed by applying a mask (light-yellow, top row) around the top ring and using particle images where the signal from the bottom ring had been subtracted. Orthogonal views of the unsharpened maps obtained from these refinements (center row), and top views of these cryo-EM maps after sharpening (bottom row).

**a**

5' CAGCCAACTAAACTA 3'
3' GTCGGTTGATTTGAT 5'

**b**

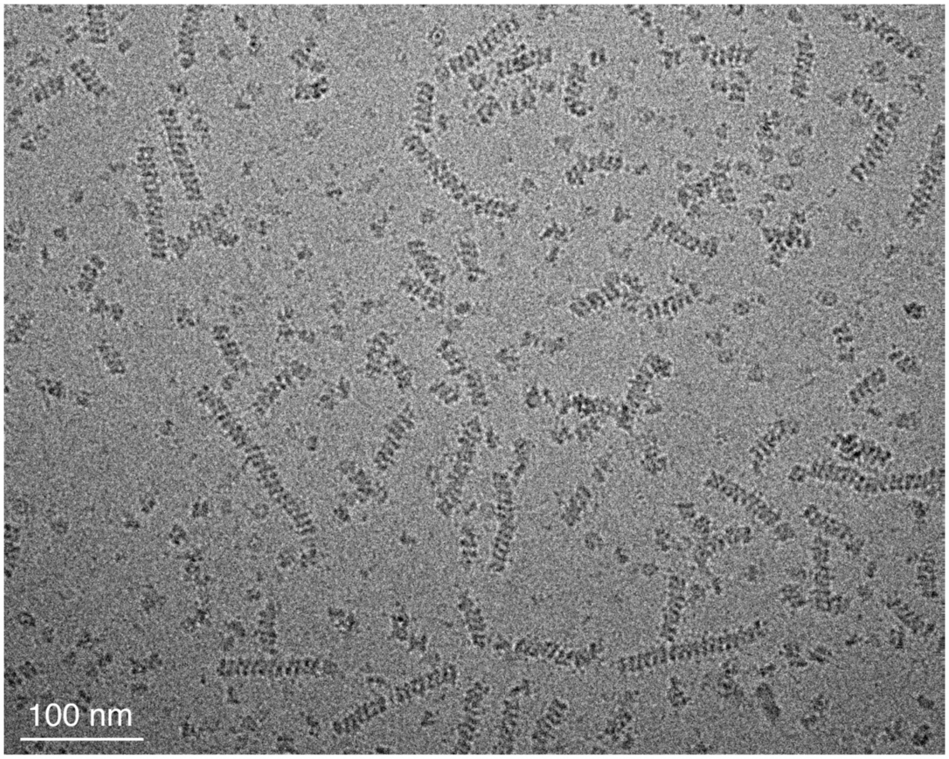


**Supplementary Fig. 8. TnsC$^{S}$-A225V rings stack end-to-end when bound to duplex DNA. (A)** Sequence of the 15 bp DNA duplex used to assemble the TnsC$^{S}$-A225V:dsDNA complexes. **(B)** Representative cryo-EM micrograph. The TnsC-DNA complexes tend to form ring stacks of different lengths.

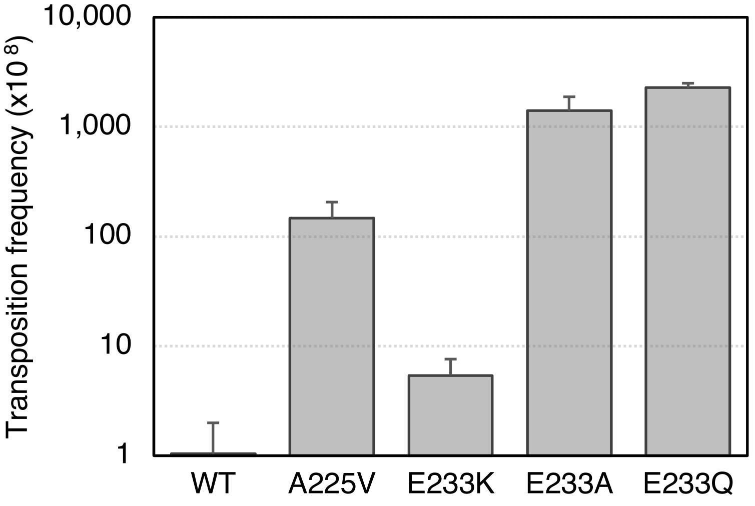


**Supplementary Fig. 9. TnsABC transposition frequency with TnsC gain-of-function mutants.** Transposition frequency for TnsABC with TnsC-A225V, TnsC-E233K, TnsC-E233A, and TnsC-E233Q measured by the lambda hop assay. Frequency is indicated as transposition events/pfu donor phage x $10^{8,}$ and the average of three experiments is indicated +/- one standard deviation (n=3).

---

**Supplementary Movie 1. Conformational rearrangements of the C-terminal region of TnsC$^{S}$-A225V.** In the conformation seen in the crystal structure (first frame), TnsC$^{S}$-A225V would not be able to form a ring because the C-terminal region of TnsC (residues 381-487, colored salmon) would overlap with the oligomerization interface. Therefore, to assemble the ring, this region must swivel ~ 180° around residue Asp402 to orient the C-terminal region outwards of the ring (last frame). The N-terminal extension (TnsD-interaction region) preceding the AAA+ domain and INS1 are colored in blue and teal for reference.
